# Structural Basis for Allosteric Control of the SERCA-Phospholamban Membrane Complex by Ca^2+^ and cAMP-dependent Phosphorylation

**DOI:** 10.1101/2020.08.28.271940

**Authors:** Daniel K. Weber, Máximo Sanz-Hernández, U. Venkateswara Reddy, Songlin Wang, Erik K. Larsen, Tata Gopinath, Martin Gustavsson, Razvan L. Cornea, David D. Thomas, Alfonso De Simone, Gianluigi Veglia

## Abstract

Phospholamban (PLN) is a mini-membrane protein that directly controls the cardiac Ca^2+^-transport response to β-adrenergic stimulation, thus modulating cardiac output during the fight- or-flight response. In the sarcoplasmic reticulum membrane, PLN binds to the sarco(endo)plasmic reticulum Ca^2+^-ATPase (SERCA), keeping this enzyme’s function within a narrow physiological window. PLN phosphorylation by cAMP-dependent protein kinase A or increase in Ca^2+^ concentration reverses the inhibitory effects through an unknown mechanism. Using oriented-sample solid-state NMR spectroscopy and replica-averaged NMR-restrained structural refinement, we reveal that phosphorylation of PLN’s cytoplasmic regulatory domain signals the disruption of several inhibitory contacts at the transmembrane binding interface of the SERCA-PLN complex that are propagated to the enzyme’s active site, augmenting Ca^2+^ transport. Our findings address long-standing questions about SERCA regulation, epitomizing a signal transduction mechanism operated by posttranslationally-modified bitopic membrane proteins.

## INTRODUCTION

Miniproteins are translated from small open reading frames of 100-300 nucleotides in length and constitute a neglected portion of the human proteome^1^. Most miniproteins are membrane-embedded and act as regulators or ancillary proteins to enzymes or receptors^2–4^. Among the most critical miniproteins is phospholamban (PLN), a bitopic membrane polypeptide that regulates the function of the sarco(endo)plasmic reticulum Ca^2+^-ATPase (SERCA) in cardiac muscle^5^. PLN directly controls cardiac output by maintaining SERCA’s activity within a tight physiological window^6^. SERCA is a ten-transmembrane (TM) pump that promotes diastole by removing Ca^2+^ from the sarcoplasm and restoring high Ca^2+^ concentrations in the sarcoplasmic reticulum (SR) in preparation for the next systole^6^. As with other P-type ATPases, SERCA is fueled by ATP and cycles between two major conformational states *E*1 and *E*2, of high and low Ca^2+^-affinity, respectively^7^. In cardiomyocytes, PLN is expressed in 4-fold molar excess of SERCA, suggesting that this endogenous regulator is permanently bound to the enzyme in a 1:1 stoichiometric ratio^8^. PLN binds the ATPase *via* intramembrane protein-protein interactions, lowering its apparent Ca^2+^ affinity and stabilizing the *E*2 state of the pump^6,9^. SERCA/PLN inhibitory interactions are relieved upon β-adrenergic stimulation, which unleashes cAMP-dependent protein kinase A (PKA) to phosphorylate PLN’s cytoplasmic domain at Ser16, enhancing Ca^2+^ transport by SERCA and augmenting heart muscle contractility^10^. Ablation, point mutations, or truncations of PLN have been linked to congenital heart disease^9^. Despite multiple crystal structures of SERCA alone^7^ and several structural studies of PLN free and bound to SERCA^11–13^, the inhibitory mechanisms of PLN and its reversal upon phosphorylation or Ca^2+^ increase are still unknown. Mutagenesis data and molecular modeling suggested that SERCA regulation occurs through electrostatic and hydrophobic interactions between the helical transmembrane (TM) region of PLN and the binding groove of the ATPase formed by TM2, TM6, and TM9^14^. Upon phosphorylation, or binding SERCA, however, the helical TM domain of PLN does not undergo significant changes in secondary structure^15–17^. As a result, X-ray crystallography^16^ and other structural techniques (*e.g.*, EPR or NMR) have not offered significant mechanistic insights into the regulatory process.

Here, we reveal the elusive signal transduction mechanism responsible for phosphorylation-induced activation of the SERCA/PLN complex using a combination of oriented-sample solid-state NMR (OS-ssNMR) spectroscopy and dynamic structural refinement by replica-averaged orientational-restrained molecular dynamics simulations (RAOR-MD)^18,19^. The analysis of anisotropic ^15^N chemical shifts (CSs) and ^15^N-^1^H dipolar couplings (DCs) of PLN alone and in complex with SERCA in magnetically aligned lipid bicelles unveiled collective topological changes of PLN’s inhibitory TM domain in response to Ser16 phosphorylation. Specifically, the local perturbations of phosphorylation were alloster-ically transmitted via an order-disorder transition of the juxtamembrane helical residues involved in several inhibitory interactions with SERCA. This intramembrane regulatory mechanism represents a potential paradigm of the structural basis of SERCA activity modulation by other regulins (*e.g.*, sarcolipin, myoregulin, DWORF, etc.)^2^ in response to different physiological cues.

## RESULTS

### The TM domain of PLN undergoes a topological two-state equilibrium

In lipid membranes, PLN adopts an *L*-shaped conformation, with a membrane-adsorbed, amphipathic regulatory region (domain Ia, M1 to T17) connected by a short loop (Ile18 to Gln22) to a helical inhibitory region (domains Ib, Gln23 to Asn30; and domain II, Leu31 to Leu52), which crosses the SR membrane^20,21^. In its storage form, PLN is pentameric^21–23^ and de-oligomerizes into active *L-shape* monomers^20^. The dynamic cytoplasmic region undergoes an order-disorder transition between tense (*T*) and relaxed (*R*) states, with the latter promoted by Ser16 phosphorylation^15,17,24^. Upon binding SERCA, domain Ia transitions to a more-rigid and non-inhibitory bound (*B*) state, becoming more populated upon phosphorylation^15,17^. How does Ser16 phosphorylation signal the reversal of inhibition to the TM region? Since the inhibitory TM region is ~ 45 Å away from Ser16 and ~ 20 Å from the SERCA’s Ca^2+^ binding sites, we speculated that both phosphorylation (of PLN) and Ca^2+^ binding (to SERCA) must transmit conformational and topological changes across the membrane, thus allosterically modulating SERCA’s function.

Residue-specific anisotropic NMR parameters such as CSs and DCs are exquisitely suited to describe topological transitions such as tilt, bend, and torque of TM proteins in lipid bilayers near-physiological conditions^25,26^. Their analysis by OS-ssNMR requires that membrane-embedded proteins are uniformly oriented relative to the static magnetic field (**B_0_**). Therefore, we reconstituted PLN free and in complex with SERCA into magnetically aligned lipid bicelles^27^. Since monomeric PLN is the functional form^28^, we utilized a monomeric mutant of PLN devoid of TM cysteine residues^29^. Both unphosphorylated (PLN^AFA^) and phosphorylated (pPLN^AFA^) variants of PLN were expressed recombinantly, while SERCA was purified from mammalian tissues^30^. Since lipid bicelles orient spontaneously with the normal of the membrane 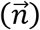 perpendicular to **B_0_**, we doped the sample with Yb^3+^ ions to change the magnetic susceptibility and orient the lipid membranes with 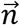 parallel to **B_0_**. This expedient doubles the values of CSs and DCs and increases the resolution of the NMR spectra^31,32^. Fig. 1a shows the 2D [^15^N-^1^H] sensitivity-enhanced (SE)-SAMPI4^33^ separated local field (SLF) spectra of free PLN^AFA^ and pPLN^AFA^. Due to PLN’s intrinsic conformational dynamics, the SLF spectra visualize only its TM region. The spectra display the typical wheel-like pattern diagnostic of a helical conformation for both the TM domains of PLN^AFA^ and pPLN^AFA^. Residue-specific assignments were carried out on free PLN^AFA^ using a combination of a 3D SE-SAMPI4-PDSD spectrum^34^, selective ^15^N labeled samples, and predictions from MD simulations^35^ (Extended Data Table S1; Extended Data Fig. 1-3). To obtain PLN’s topology in lipid bilayer, the assigned resonances were fit to idealized *P*olar *I*ndex *S*lant *A*ngle (PISA) models extracting whole-body tilt (θ) and rotation or azimuthal (ρ) angles^36,37^, which for free PLN^AFA^ was θ = 37.5 ± 0.7° and ρ_L31_ = 201 ± 4°, and for pPLN^AFA^ θ = 34.8 ± 0.5° and ρ_L31_ = 201 ± 4°, where ρ_L31_ is the rotation angle referenced to Leu31. Notably, the high resolution of the oriented SLF spectra of PLN^AFA^ show two distinct sets of peaks (Fig. 1b), with populations unevenly distributed. The average population of the minor state estimated from the normalized peak intensities is approximately 30 ± 9 %. Remarkably, the resonances of the minor population overlap almost entirely with those of pPLN^AFA^ (Fig. 1b), revealing a topological equilibrium in which the TM region of PLN interconverts between two energetically different orientations. We previously showed that PLN phosphorylation shifts the conformational equilibrium toward the *R* state^17^, releasing the interactions with the lipid membranes of domain Ia (Fig. 1c). Our OS-ssNMR data show that these phosphorylation-induced effects propagate to the TM domains, shifting the topological equilibrium toward the less populated state.

**Fig. 1:**
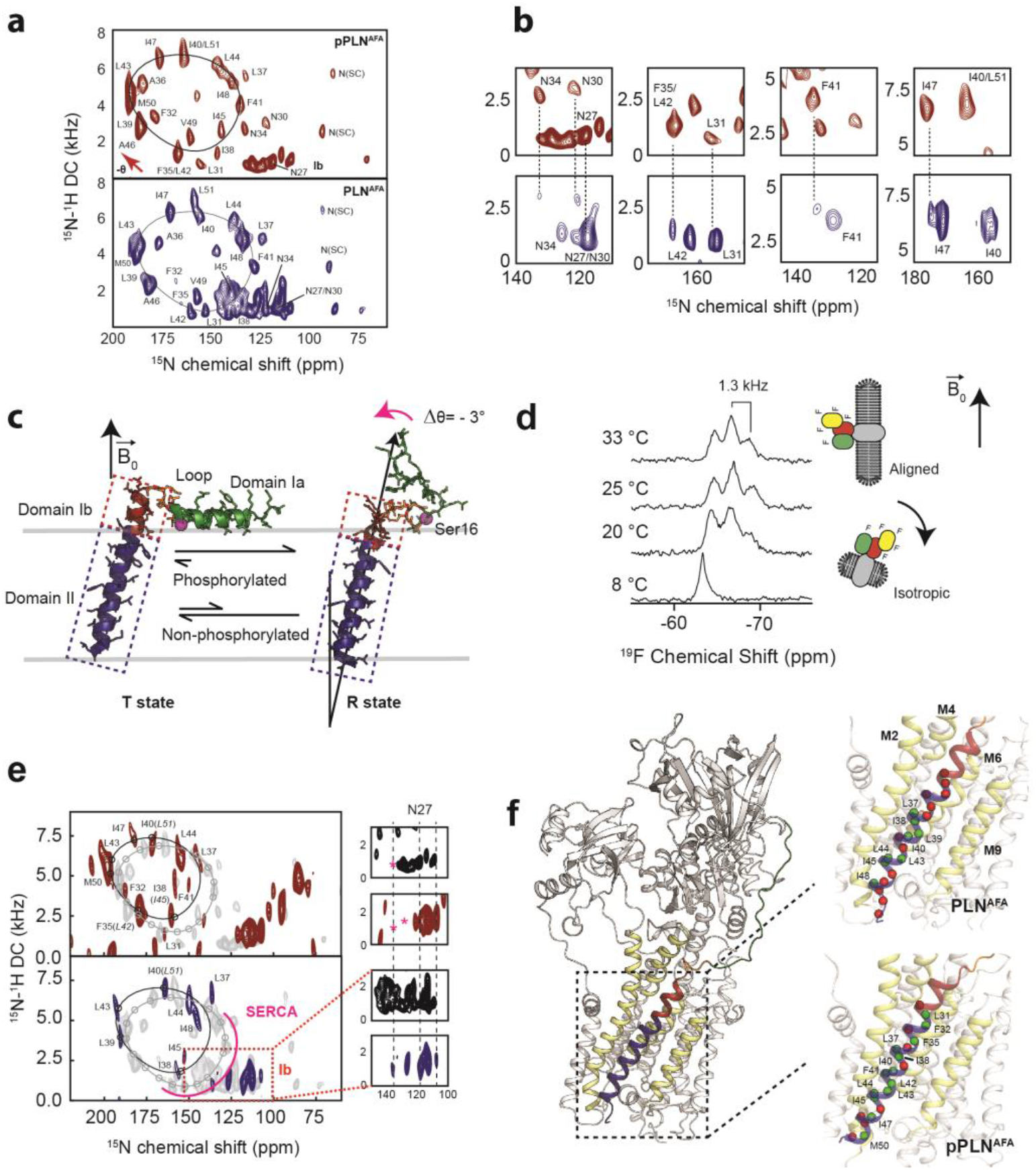
Topological equilibrium of PLN and pPLN free and bound to SERCA in lipid bilayers detected by OS-ssNMR. **a,** 2D [^15^N-^1^H] SE-SAMPI4 spectra of PLN^AFA^ and pPLN^AFA^ reconstituted into aligned lipid bicelles. The fitting of resonance patterns with PISA wheels for an ideal helix [(ϕ, ψ)=(−63°, −42°)] is superimposed. **b,** Expanded regions of PLN^AFA^ ^15^N-labeled at N, L, F, or I residues (lower panel, blue contours) showing two populations. The upper panels (red) are the corresponding regions for the U-^15^N labeled pPLN^AFA^. U-^15^N labelled spectra were acquired at higher signal-to-noise to observe the second population. **c,** Structures of the *T* (PDB 2KB7^20^) and *R* (PDB 2LPF^41^) states for PLN^AFA^. **d,**^19^F NMR spectra of TFMB-tagged SERCA reconstituted into anisotropic (*q* = 4) bicelles at variable temperatures. **e,** 2D [^15^N-^1^H] SE-SAMPI4 spectra of uniformly ^15^N labeled PLN^AFA^ (blue, lower panel) and pPLN^AFA^ (red, upper panel) bound to SERCA in the absence of Ca^2+^ (*E*2 state). Spectra are overlaid with PLN^AFA^ or pPLN^AFA^ in their free forms (grey). PISA wheels are overlaid, showing assigned residues (black points) used to fit helical tilt and rotation angles. Ambiguous assignments are shown in parentheses. The region corresponding to domain Ib is expanded to show peak broadening (asterisk) following the transition of PLN’s cytoplasmic region to the *B* state. **f,** Selected structure of the SERCA/PLN^AFA^ complex. Expanded region shows visible (green spheres, labelled) and broadened (red spheres) residues mapped onto domain II of PLN^AFA^ (left) and pPLN^AFA^ (right, form a structure of the SERCA/PLN^AFA^ complex).

### PLN phosphorylation by PKA signals a rearrangement of the SERCA/PLN binding interface

To investigate how Ser16 phosphorylation allosterically affects the inhibitory TM binding interface, we reconstituted the SERCA/PLN complex in lipid bicelles and studied it by OS-ssNMR. The alignment of mammalian SERCA in bicelles was confirmed by cross-linking the most reactive cysteines with a trifluoromethylbenzyl (TFMB)-methanethiosulfonate (MTS) tag to probe its alignment by ^19^F NMR (Fig. 1d and Extended Data Fig. 4). Five of the twenty-four cysteines of SERCA were uniquely labeled as monitored by solution NMR in isotropic bicelles (*q* = 0.5). In anisotropic bicelles (*q* = 4.0) and at low temperature, the ssNMR spectrum of ^19^F-SERCA consists of a single unresolved ^19^F resonance due to the rapid reorientation of the enzyme in the isotropic phase. Upon increasing the temperature, the ^19^F-SERCA/bi-celle complex orients with 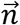 perpendicular to **B**_0_, and the ^19^F resonance becomes anisotropic as a triplet with 1.3 kHz dipolar coupling^38^. Fig. 1e shows the 2D SLF spectra of PLN^AFA^ and pPLN^AFA^ in complex with SERCA. To maintain a functional and stable complex, we used a lipid-to-complex molar ratio of 2000:1, with PLN concentration 10 times less than in the SERCA free samples. Therefore, the signal-to-noise ratio in the oriented spectra is significantly reduced relative to the free forms. Nonetheless, the SLF spectra of both SERCA/PLN^AFA^ and SERCA/pPLN^AFA^ complexes show the wheel-like patterns typical of the α-helical domains with selective exchange broadening for resonances located at the protein-protein binding interface (Fig. 1f). The assigned peaks associated with the helical domain II were fit to the ideal PISA model, yielding θ = 33.2 ± 1.2° and ρ_L31_ = 193 ± 7°. Therefore, upon binding SERCA, PLN^AFA^ requires a distinguishable −4.3 ± 1.4° change in tilt and a less significant −8 ± 8° change in rotation. These error bounds factor the linewidths and variation associated with substituting ambiguous assignments into the PISA fitting (parentheses of Fig. 1e). Similarly, the PISA model for pPLN^AFA^ was fit to θ = 30.4 ± 1.1° and ρ_L31_ of 197 ± 4°, suggesting that the topology of the TM domain requires adjustments of −3.4 ± 1.2° and −4 ± 6° to form a complex with the ATPase.

Reductions in the TM helix tilt angle, which accompanied phosphorylation and complex formation, also coincided with a dramatic broadening of peaks in the cluster of isotropic resonances around 140 ppm (Fig. 1e). These resonances are attributed to the dynamic domain Ib residues, and their disappearance is consistent with SERCA binding, which requires the unwinding of the juxtamembrane region, and a concomitant reduction of the tilt angle to re-establish hydrophobic matching with the thickness of the lipid bilayer^13,17^. Therefore, for free and bound PLN, we find that communication between cytoplasmic and intramembrane environments is transduced via domain Ib dynamics. Analysis of the SLF spectra also shows that phosphorylation of PLN at Ser16 restores the intensities of most resonances except for those at the upper binding interface (*i.e.*, Asn30, Leu31, Asn34, and Phe41). These spectral changes suggest a reorganization of PLN-SERCA packing interactions, rather than a complete dissociation of the complex, consistent with prior MAS-ssNMR, EPR, and FRET measurements^13,17,39,40^.

### Dynamic structural refinement of the SERCA-PLN complexes

To determine the structural ensembles of the SERCA/PLN^AFA^ and SERCA/pPLN^AFA^ complexes, we incorporated the data from our experimental measurements into RAOR-MD samplings^18,19^. This dynamic refinement methodology employs full atomic MD simulations in explicit lipid membranes and water and utilizes restraints from sparse datasets to generate experimentally-driven structural ensembles. As starting coordinates for our samplings, we used the X-ray structure of *E*2-SERCA/PLN, where a super-inhibitory mutant of PLN was used to stabilize the complex for crystallization^16^. We docked the TM domains of PLN^AFA^ using restraints obtained from chemical cross-linking experiments for both cytoplasmic and luminal sites^14,42,43^ (Extended Data Fig. 5a). The dynamic cytoplasmic region (loop and domain Ia), which was not resolved in the crystal structure, was held in proximity to the nucleotide-binding (N) and phosphorylation (P) domains of SERCA using upper boundary restraints from paramagnetic relaxation enhancements (PRE) obtained from MAS-ssNMR^17^. Additionally, CSs and DCs from OS-ssNMR were applied to the TM region as ensemble-averaged restraints across eight replicas. The resulting structural ensembles were in excellent agreement with all the available experimental data for both complexes. The overall profile of the average pairwise distances for residues in domain Ia and loop to the spin-label at Cys674 matches the PRE measurements, with the minimal distance (*i.e.,* maximum PRE effect) observed for PLN-Tyr6 (Extended Data Fig. 5b,c). Similarly, back-calculated CS and DC values for PLN^AFA^ and pPLN^AFA^ were in excellent agreement with experiments (Extended Data Fig. 5d). Average back-calculated tilt angles of 32.8° and 30.4° for PLN^AFA^ and pPLN^AFA^, respectively, matched PISA fits to experimental values (Extended Data Fig. 5e,6). All pairwise distances between previously reported cross-linkable positions^14,42–44^ were distributed within acceptable ranges (Extended Data Fig. 5f). Although not used as an initial docking restraint, cytoplasmic residues PLN Lys3 and SERCA Lys397 were also partially distributed within a distance consistent with previously reported cross-linking^45^.

To assess the SERCA/PLN complexes’ conformational landscape, we used principal component analysis (PCA). PCA identified a combined opening of the cytoplasmic headpiece involving a hinge-like displacement of the N domain and rotation of the A domain away from the P domain (PC1) and planar rotations separating the N and A domains (PC2) (Fig. 2a,b, Extended Data Fig. 7, Extended Data Movie 1). These motions differentiate the *E*1 and *E*2 states of SERCA, as shown by the projections of crystal structures onto the PCA map (Fig. 2c,d). The SERCA/PLN^AFA^ complex spans four distinct clusters, while the SERCA/pPLN^AFA^ complex spans eight (see Extended Data Fig. 8 for representative structures). When bound to PLN^AFA^, SERCA mostly retains the compact *E*1-like headpiece present in the crystal structure and interconverts with equal frequency between a highly compact cluster (3) resembling nucleotide-bound *E*1 states and a cluster (1) intermediate toward the *E*2 states, which exhibits a partial opening of the A and N domains caused by breaking of the salt bridges involving Arg139-Asp426/Glu435 and Lys218-Asp422. Albeit biased along the *E*1 coordinate of PC2, similar states were present for pPLN^AFA^, but the interaction of pSer16 with Arg604 weakens the Asp601-Thr357 and Arg604-Leu356 hydrogen bonds at the hinge of the N and P domains, leading to four additional open states (clusters 5 to 8). Separate clusters correspond to successive breakages of interdomain hydrogen bonds in the headpiece. Salt bridges between PLN-Ser16 to SERCA-Arg460 and PLN-Arg14 to SERCA-Glu392 were also found to stabilize these open states (Extended Data Fig. 9). These open states resemble off-pathway crystal structures solved for the Ca_2_*E*1 state observed in the absence of nucleotide^7,46,47^. For all clusters, the binding interactions near the Ser16 position were more persistent for pPLN^AFA^ than PLN^AFA^ (Fig. 2e,f, Extended Data Fig. 8).

**Fig. 2:**
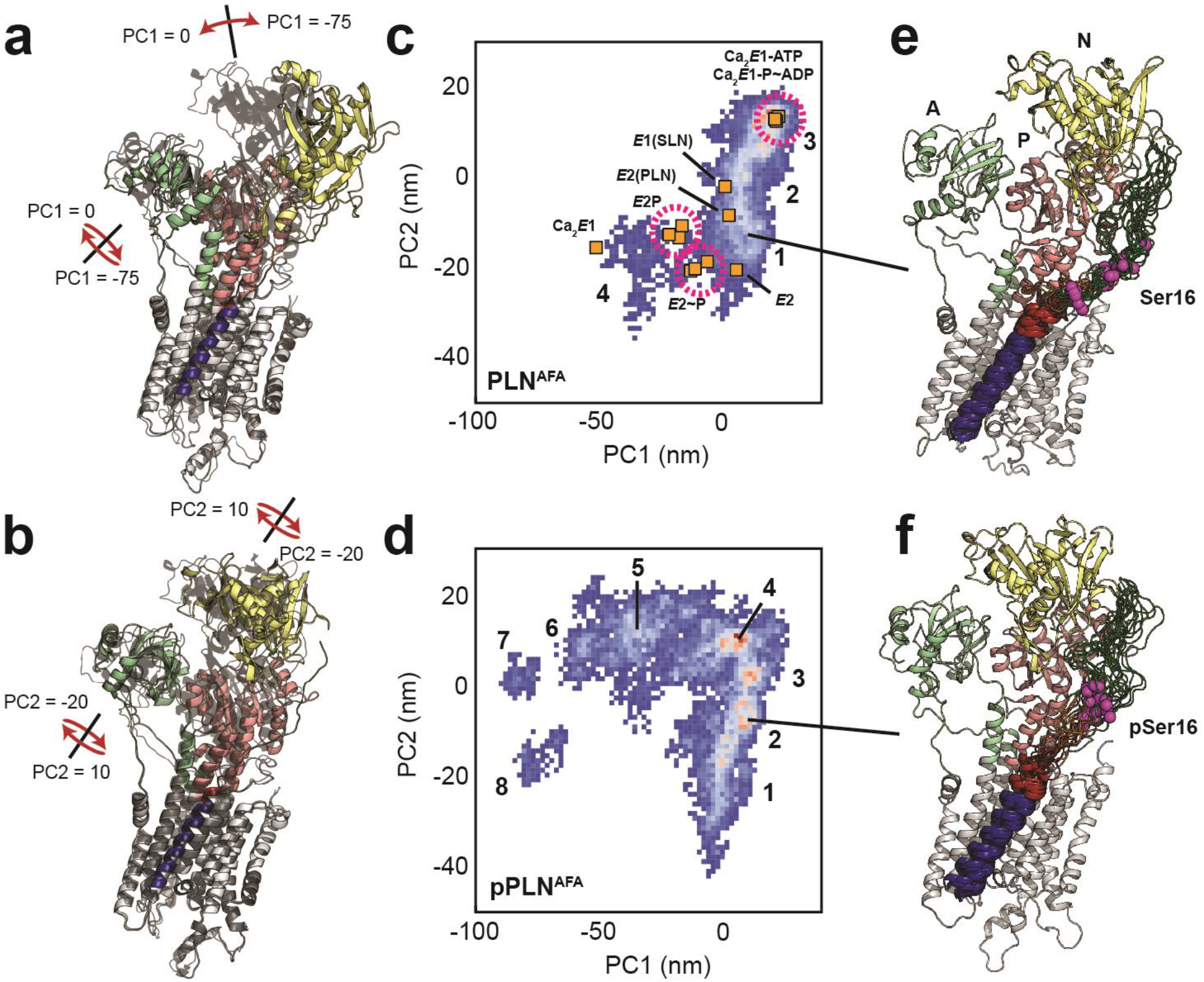
Conformational ensembles and energy landscapes of the SERCA/PLN^AFA^ and SERCA/pPLN^AFA^ complexes. **a, b,** Depiction of headpiece movements associated with the first (**a**) and second (**b**) principal components. Structures with the highest PC values are shown as transparent black. **c, d,** PCA histograms of SERCA/PLN^AFA^ (**c**) and SERCA/pPLN^AFA^ (**d**) structural ensembles with projections of crystal structures in various states: Ca_2_*E*1-ATP^48,49^, Ca_2_*E*1~P-ADP^49–51^, Ca_2_*E*1^46^, *E*1-SLN^52^, *E*2-PLN^16^, *E*2P^50,53^, *E*2~P^50,53,54^, and *E*2^55^. Clusters are numbered. **e,** f, Top 20 most representative structures of PLN^AFA^ (**e**, cluster 1) and pPLN^AFA^ (**f**, cluster 2) bound to SERCA from the most representative state.

From the analysis of the structural ensembles of the two complexes, it emerges that the relief of inhibition (i.e., activation) occurs via a rearrangement of the intramembrane contacts between the TM region of pPLN^AFA^ and SERCA, with a reconfiguration of electrostatic interactions near the phosphorylation site and a disruption of packing at the protein-protein interface (Fig. 3a,b). The interactions between the cytoplasmic regions are transient and highly dynamic, resembling the conformational ensembles of intrinsically disordered complexes^56^. For both complexes, we observe persistent interactions between PLN-Glu2 and SERCA-Lys365, PLN-Glu19, and SERCA-Lys328, and to a lesser extent PLN-Lys3 and SERCA-Asp399 and PLN-Tyr6 and SERCA-Asp557. For PLN^AFA^, however, the arginine residues (Arg9, Arg13, and Arg14) interact transiently with SERCA-Glu606, while for pPLN^AFA^, the phosphate group at Ser16 interacts strongly with SERCA-Arg604 and SERCA-Lys605. Interestingly, we detected the formation of intramolecular salt-bridges between the phosphate of Ser16 and PLN-Arg9, PLN-Arg13, and PLN-Arg14, causing domain Ia to adopt a compact conformation as previously suggested by fluorescence data^57,58^ (Extended Data Table 2, 3). These cytoplasmic protein-protein interactions destabilized PLN’s domain Ib and consequently severed inhibitory intermolecular contacts with SERCA’s TM helices involving Gln23-Leu321/Arg325 (M4), Lys27-Phe809 (M6) and Asn30-Trp107 (M2), while the intermolecular contacts involving domain II are mostly retained (Fig. 3a and Extended Data Fig. 10). In fact, hydrophobic substitutions within domain Ib have been identified as hotspots for engineering super-inhibitory PLN mutants (*i.e.*, Asn/Lys27Ala and Asn30Cys), which exhibit stable helical structure well into the loop domain^16,59,60^. Importantly, these structural ensembles capture the order-disorder dynamics of domain Ib resonances observed in the OS-ssNMR spectra and suggested by previous NMR and EPR studies^61^. Destabilization and detachment of this region is consistent with the reappearance of exchange-broadened interfacial resonances of domain II paralleled by broadening of the resonances of the dynamic domain Ib in the SLF spectra of the SERCA/pPLN^AFA^ complex (Fig 1e). The interactions of domain Ia (Arg13-Ser16) and inhibitory contacts of domain Ib (at Lys27) with SERCA appear to be mutually exclusive (Fig. 3c-f). In both complexes, the electrostatic interactions of PLN-Arg13, PLN-Arg14, or PLN-pSer16 with the SERCA’s Arg604-Glu606 stretch cause the detachment of PLN’s domain Ib and the consequent weakening of the inhibitory interaction (Fig. 3c-g). This illustrates the regulatory role of the *B* state of PLN for relieving inhibition and the superinhibitory activity of domain Ia-truncated PLN^17^.

**Fig. 3.**
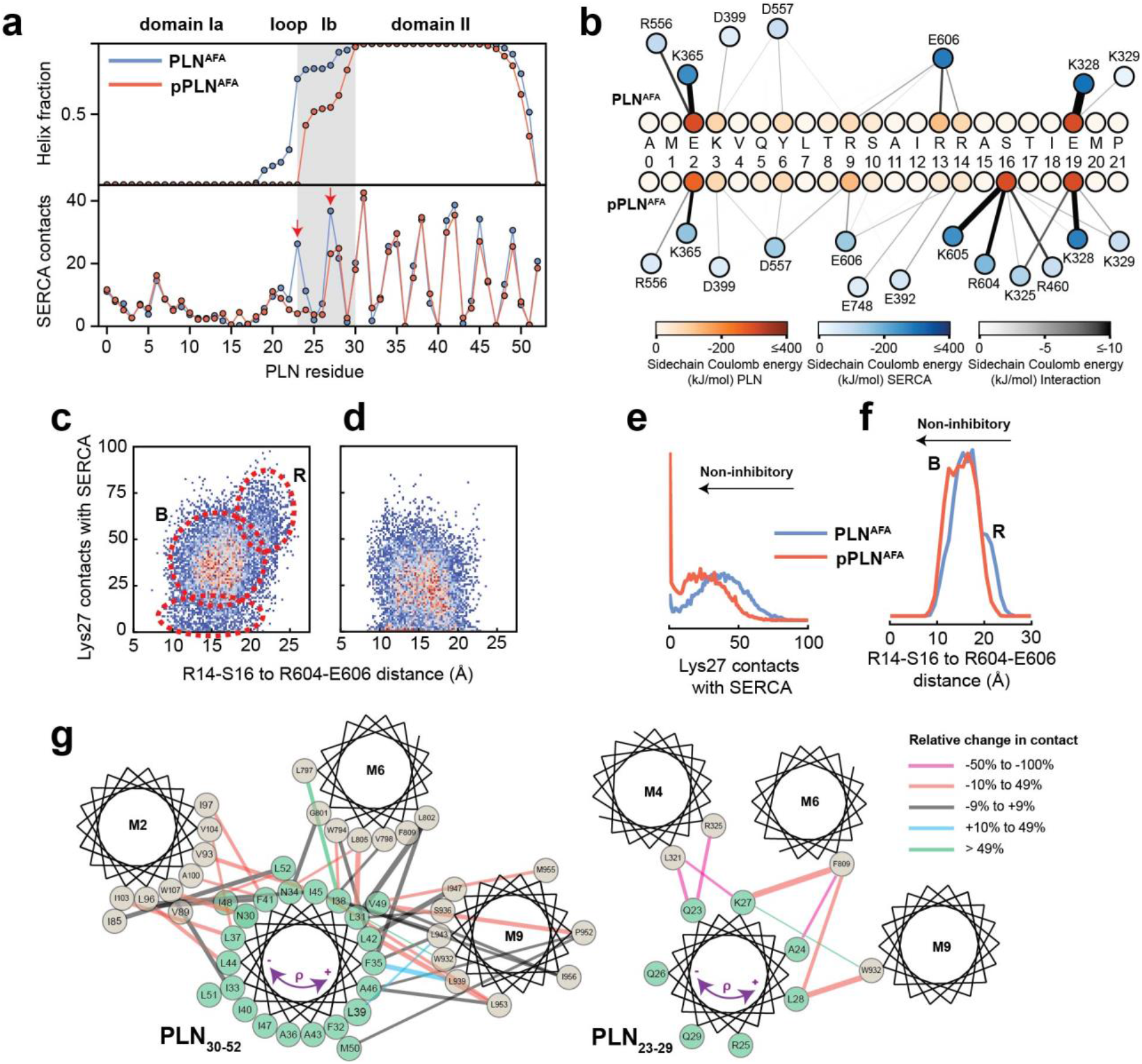
Mechanism for reversal of PLN inhibition by phosphorylation. **a,** Ensemble-averaged per-residue structural analysis (upper panel) and intermolecular contact profiles (lower panel). A contact is defined when any PLN atom comes within 3.5 Å of any SERCA atom for any given frame. **b,** Spider plot of pairwise electrostatic interactions between cytoplasmic residues and SERCA in RAOR-MD conformational ensembles. **c, d,** 2D Histograms correlating the distances between the cytoplasmic binding interfaces, defined by the center of masses of Arg14 to Ser16 of PLN and Arg604 to Glu606 of SERCA, to the inhibitory intermolecular contacts of PLN Lys27 for the SERCA/PLN^AFA^ (**c**) and SERCA/pPLN^AFA^ (**d**) ensembles. **e, f,** Corresponding 1D histograms for Lys27 contacts (**e**) and binding of the cytoplasmic domain (**f**). **g.** Disruption of the inhibitory TM pairwise interactions detected in the RAOR-MD conformational ensembles. Linewidths of the interhelical contacts are scaled to average contacts per frame for the non-phosphorylated complex and colored by the relative change observed with phosphorylation. Directions of the purple arrows exemplify clockwise or counterclockwise rotations of the TM domain of PLN during the trajectories.

### Phosphorylation disrupts correlated motions between PLN^AFA^ and SERCA’s Ca^2+^ binding sites

To assess the effects of PLN’s phosphorylation on Ca^2+^ transport, we calculated the topological correlations of the TM’s tilt angle fluctuations between PLN’s and SERCA’s TM helices (Fig. 4a-d). When PLN^AFA^ is bound to SERCA, we observe a dense network of correlated motions between PLN’s TM region and the binding groove (TM2, TM6, and TM9), as well as a dense cluster of correlations involving TM3, TM4, TM5, TM6, and TM7. This allosteric coupling influences the Ca^2+^ binding sites’ geometry, possibly reducing SERCA’s Ca^2+^ binding affinity. In contrast, the analysis of the trajectories of the SERCA/pPLN^AFA^ complex displays only correlated motions between the TM of pPLN^AFA^ and the most proximal SERCA helices, with only a sparse network of correlations involving TM4, TM5, TM6, and TM8. Phosphorylation of PLN at Ser16 increases the electrostatic interactions with the cytoplasmic domain of SERCA (*R* to *B* state transition, *i.e.*, disorder to order)^17^, and simultaneously weakens intramembrane protein-protein interactions to uncouple the dynamic transitions of PLN from SERCA. The latter removes the structural hindrance of PLN’s TM domain and augments Ca^2+^ transport (Fig. 4e,f).

**Fig. 4.**
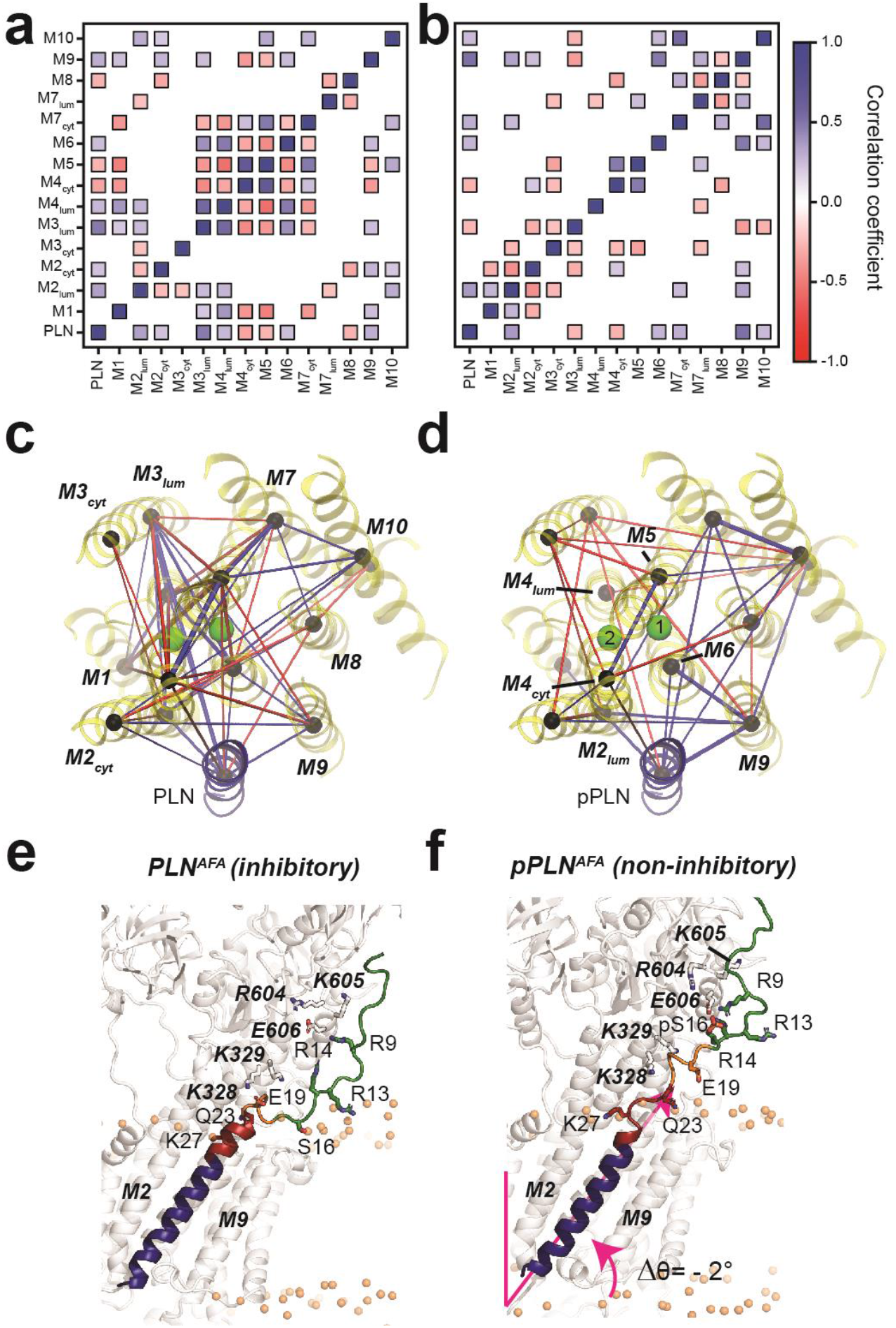
PLN topological transitions are allosterically coupled to SERCA’s Ca^2+^ binding sites. **a, b,** Correlation maps of motions between the TM topology of PLN^AFA^ (**a**) or pPLN^AFA^ (**b**) with the topology of the 10 TM domains of SERCA. **c, d,** Corresponding spider plots showing the density of correlations are displayed below. Green spheres mark positions of the calcium-binding sites. **e, f,** Snapshots of the SERCA/PLN^AFA^ (**e**) and SERCA/pPLN^AFA^ (**f**) complexes highlighting the transient interactions with the cytoplasmic region and loosened interactions with the TM region of SERCA.

### Effects of Ca^2+^ ion binding to SERCA on PLN’s topology

To assess the effects of Ca^2+^, we performed SLF experiments on SERCA/PLN^AFA^ and SERCA/pPLN^AFA^ complexes in the *E*1 state (Extended Data Fig. 11a-c). The addition of Ca^2+^ to the SERCA/PLN^AFA^ complex did not cause significant changes to the PLN topology (θ = 32.9 ± 1.4° and ρ_L31_ = 199 ± 4°). However, the reappearance (*i.e.*, sharpening) of several resonances in the spectra (*e.g.*, Phe32, Phe35/Leu42, and Ala36) indicates a rearrangement of the binding interface between the two proteins similar to phosphorylation’s effect on the *E*2 complex. A small topological change, however, was observed for the SERCA/pPLN^AFA^ complex, for which Ca^2+^-binding induced a decrease of both tilt and rotational angles of 1.8 ± 1.3° and 10 ± 7°, respectively (θ = 28.6 ± 0.7° and ρ_L31_ = 187 ± 6°). Due to the lack of X-ray structures, we were unable to carry out dynamic modeling of these complexes. However, these experimental results agree with our *E*2-SERCA models suggesting that a loss of inhibition, either from phosphorylation or Ca^2+^ binding, does not require an extensive structural and topological reconfiguration of PLN’s domain II or complete dissociation of the complex.

## DISCUSSION

OS-ssNMR spectroscopy revealed that the TM helix of PLN undergoes a topological equilibrium that is shifted upon phosphorylation, providing direct evidence of the allosteric coupling between the outer membrane regulatory and TM inhibitory regions. Our dynamic modeling of the SERCA/PLN complexes using experimental restraints shows that the structural disorder of the juxtamembrane domain Ib following Ser16 phosphorylation of PLN signals a slight topological change in the TM region that is sufficient to relieve its inhibitory function. This event involves allosteric effects between the inhibitory interactions of domain II of PLN and SERCA’s core helices harboring the Ca^2+^ binding sites. In addition to the localized disruption of PLN domain Ib interactions with SERCA, phosphorylation and Ca^2+^-binding signal a collective switch of PLN’s TM domain from an inhibitory to a non-inhibitory topology. The tilt angle reductions of PLN accompanying the relief of inhibition are easily identifiable from the OS-ssNMR spectra, while rotations are more subtle; nonetheless these topological changes are sufficient to disrupt critical inhibitory interactions.

Recent X-ray investigations and extensive computational studies showed that SERCA undergoes significant rocking motions throughout its enzymatic cycle^62–64^. These conformational transitions analyzed in the absence of PLN are highly concerted and cooperative, *i.e.*, the dynamics of the cytoplasmic headpiece of SERCA correlates with its TM domains^63,64^. PLN (and other regulins)^2^ wedges into the ATPase’s binding groove and either correlate with the topological changes of SERCA’s TM domains or interfere with its rocking motions leading to uncoupling of ATP hydrolysis and Ca^2+^ transport. PLN experiences significant changes in tilt angle (ranging from 28.6° to 37.5°), depending on both the conformational state of PLN and enzymatic state of SERCA. Therefore, these dynamic and topological transitions provide the mechanism to modulate TM protein-protein interactions, which can be tuned by posttranslational phosphorylation, O-glycosylation^65^, and binding of ancillary proteins^66,67^.

The structural and dynamic changes of PLN, detected by OS-ssNMR, resolves an ongoing controversy about the *subunit* vs. *dissociative* models proposed for SERCA regulation^9,39,40,68,69^. The latter model speculates that the reversal of the inhibitory function of PLN is due to a complete dissociation of this regulin from the ATPase; but this is not supported by spectroscopic data either *in vitro* or *in cell*^13,17,39,40,69^. On the other hand, the subunit model agrees well with all spectroscopic measurements, but it does not explain the reversal of inhibition caused by phosphorylation or the elevation of Ca^2+^ concentration. Our ssNMR-driven dynamic calculations clearly shows that topological and structural changes modify the interactions at the interface and are propagated to the distal Ca^2+^ binding sites.

Fig. 6 summarizes our proposed mechanistic model for allosteric control of SERCA by PLN’s topological changes. We previously showed that PLN’s cytoplasmic domain undergoes a three-state equilibrium (*T*, *R*, and *B*) in which the *T* and *R* states are inhibitory, while the *B* state is non-inhibitory^17^. Our new data show that, when bound to the *E*2*-*SERCA state, the TM region of PLN (domains Ib and II) remains locked into the ATPase’s binding groove. An increase of Ca^2+^ concentration drives SERCA into the *E*1 state and reconfigures the intramembrane binding interface to augment Ca^2+^ transport without significant topological changes to PLN. On the other hand, detectable topological changes occur upon PLN’s phosphorylation, both at low and high Ca^2+^ concentrations, as its conformational equilibrium is shifted toward the non-inhibitory *B* state^70^ and changes are transmitted across the SERCA/PLN interface to increase Ca^2+^ transport. In this framework, it is possible to explain how single-site disease mutations or deletion in domains Ia and Ib may lead to perturbations of intra- and extra-membrane protein-protein interactions that result in dysfunctional Ca^2+^ transport^59,71,72^.

**Fig. 5.**
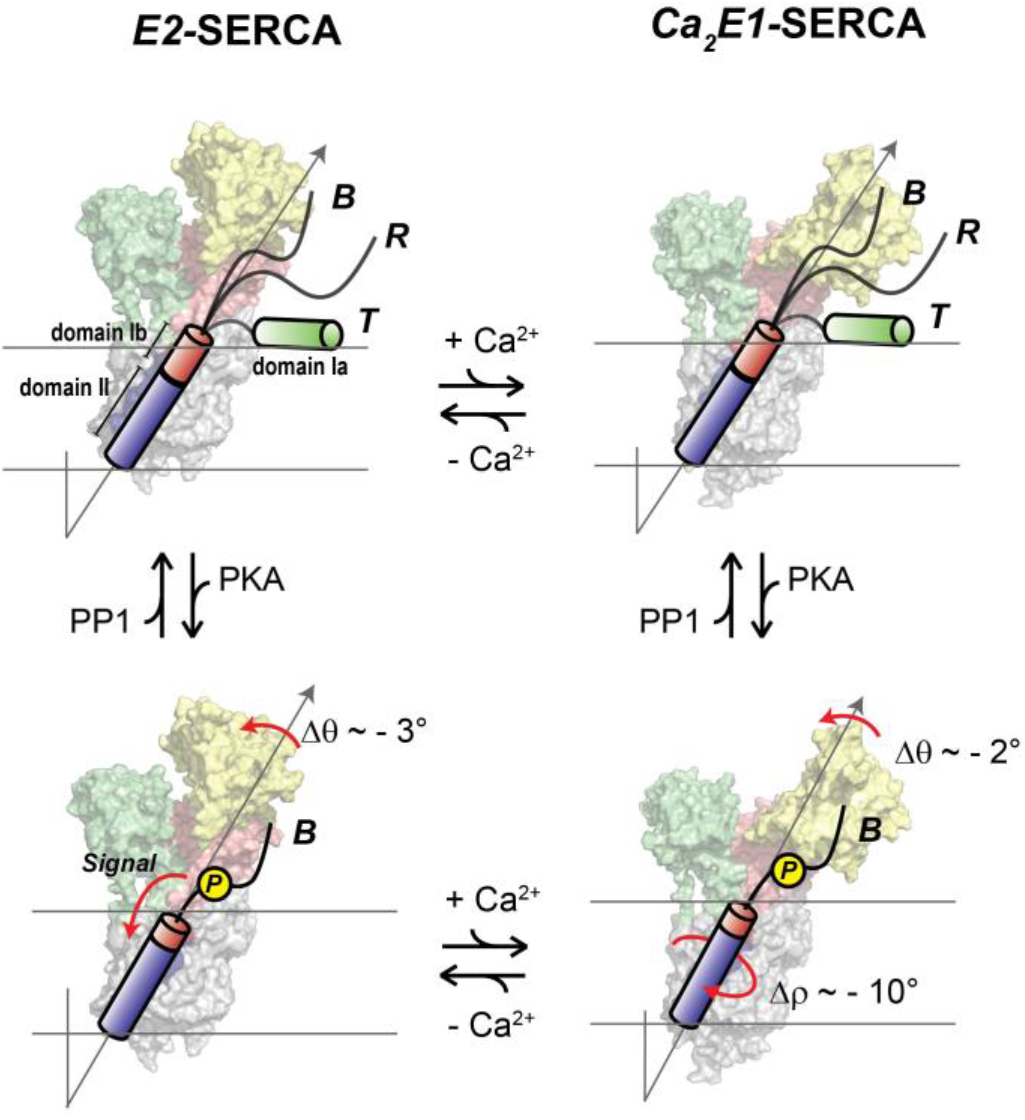
Regulatory model of SERCA by PLN’s phosphorylation and Ca^2+^. PLN’c cytoplasmic domain exists in equilibrium between three distict populations in the presence of SERCA. The increase of Ca^2+^ ions causes significant shifts SERCA’s conformation toward the E1 state, while the topology of PLN is only slightly affected (top equilibrium). Phosphorylation of PLN at Ser16 signals more extensive topological changes with a reconfiguration of SERCA/PLN TM interactions both at low and high Ca^2+^ concentrations (bottom equilibrium), augmenting Ca^2+^ transport. Note that the T and R populations are not represented for clarity.

In conclusion, the structural dynamics and topological allostery identified for PLN may explain how bitopic miniproteins, despite their simple architecture, can fulfill diverse regulatory roles and how posttranslational modification at cytoplasmic sites may constitute switches for signal transduction across cellular membranes operated by single or multiple transmembrane domains^73^. Several mini-membrane proteins regulate membrane-embedded enzymes or receptors^1^. In the heart, phospholemman^74–76^, a member of the FXYD family, regulates the Na^+^/K^+^-ATPase interacting via its transmembrane domain, with its regulatory interactions modulated by protein kinases A and C. Several regulins have also been recently found to control SERCA’s isoforms in other tissues^2^ and share similar topologies to PLN. They all bind at distal locations from the active sites (*e.g.*, ATP or ion channels) of enzymes, revealing possible hotspots for allosteric control by small molecules. Therefore, the characterization of the topological allosteric control of SERCA by PLN represents a first step in understanding how and why evolution has preserved these small polypeptides as a means to regulate the function of ATPases^75,77^ or other membrane transporters^78^.

## Acknowledgments

This work was supported by the National Institute of Health grants R01 GM064742 and R01 HL144100 to G.V. and R01HL139065 and R37AG026160 to D.D.T. and R.L.C. D.W. was supported by an American Heart Association Postdoctoral Fellowship (19POST34420009).

## Methods

### Expression, purification, and phosphorylation of PLN^AFA^

Uniformly ^15^N-labelled PLN^AFA^ was expressed as a soluble fusion with maltose binding protein (MBP) as reported previously^1^, with minor modifications. Freshly transformed *E. coli* Codon Plus (Invitrogen) cells were used to inoculate overnight LB cultures, which were subsequently centrifuged and resuspended into M9 minimal media (^15^NH_4_Cl as the sole nitrogen source) at an OD_600_ of ~0.7. Cultures were grown at 30°C to OD_600_ 1.0 then induced with 1 mM IPTG over 20 h to a final OD_600_ of 5. Cells (~6 g/L of M9) were stored at −20°C. Selectively ^15^N-labelled PLN was expressed from M9 media (free of NH_4_Cl) with 125 mg/L of the respective ^15^N-amino acid, 300 mg/L of non-scrambling and 450 mg/L scrambling-prone ^14^N-amino acids^2^. Reverse-labeled PLN was expressed in M9 minimal media (^15^NH_4_Cl) with 1 g/L of the respective ^14^N-labeled amino acid. Induction times for selective and reverse labeling growths were reduced to 3 to 4 h to reduce scrambling.

For purification, cells were homogenized (Sorvall Omni Mixer) and lysed by sonification in 200 mL lysis buffer (20 mM sodium phosphate, 120 mM NaCl, 2 mM DTT, 1 mM EDTA, 0.1 mg/mL lysozyme, 0.5 % glycerol, 0.5% Tween 20 and protease inhibitors, pH 7.3). The lysate was centrifuged (17,500 rpm, JA25.50 rotor, 4°C, 40 min) and supernatant loaded onto 30 mL bed volume of amylose resin. The resin was washed with buffer (20 mM sodium phosphate, 120 mM NaCl, pH 7.3) and eluted into 100 mL buffer including 50 mM maltose. Elution volumes were concentrated to ~50 mL and dialyzed overnight against 3 L of cleavage buffer (50 mM Tris-HCl, 2 mM β-mercaptoethanol, pH 7.3). All purification steps were done at 4°C and yielded up to 120 mg of fusion protein from 1 L M9 media.

To phosphorylate PLN^AFA^ at Ser16 (pPLN^AFA^), the MBP fusion was dialyzed into 30 mM Tris-HCl, pH 7.5, followed by addition of 11x reaction buffer to reach 50 mM Tris-HCl, 10 mM MgCl_2_, 0.05 mM PMSF, 1 mM NaN_3_, 1 mM EDTA). The catalytic subunit of protein kinase A (PKA) was added at 1:1000 ratio to fusion protein, with 2 mM DTT, and the reaction started by addition of 2 mM ATP and incubation at 30°C for 3 h with gentle agitation.

MBP-PLN, either phosphorylated or non-phosphorylated, was cleaved with TEV protease and 2 mM DTT for 3 h at 30 °C to liberate insoluble PLN (Extended Data Fig. 12a), which was pelleted by centrifugation and dissolved into 10% SDS and 50 mM DTT at approximately 10 mg/mL then stored at −20 °C. PLN was further purified by HPLC using a Vydac 214TP10154 C4 column heated at 60 °C and eluted using H_2_O/0.1% trifluoroacetic acid (TFA) and a linear gradient of isopropanol/0.1% TFA from 10 to 40% over 10 mins then to 80% over 50 min (2 mL/min flow rate). The protein was lyophilized. Complete phosphorylation of pPLN^AFA^ was confirmed by MALDI-MS (Extended Data Fig. 12b). The inhibitory activity of PLN^AFA^ against SERCA, and relieved inhibition of pPLN^AFA^, in the DMPC/POPC (4:1) lipid bilayer composition used for NMR studies, was confirmed by a coupled enzyme assay^3^ (Extended Data Fig. 12c).

### Preparation of oriented bicelle samples

Long-chain lipids DMPC (37.0 mg), POPC (10.4 mg) and PE-DTPA (0.9 mg; *i.e*., 79.25:19.75:1.0 molar ratio) were aliquoted together from chloroform stocks (Avanti Polar Lipids), dried to a film with N_2_ and residual solvent removed under high vacuum. The film was resuspended into 1 mL ddH_2_O, freeze-thawed three times between liquid N_2_ and a 40 °C water bath then lyophilized. DHPC (7.7 mg; 1:4 molar ratio, or *q* = 4, to long chain lipids) was prepared separately from a chloroform stock, dried and lyophilized from ddH_2_O.

For bicelles containing only PLN, DHPC was dissolved into 250 μL sample buffer (20 mM HEPES, 100 mM KCl, 1 mM NaN_3_, 2.5% glycerol, pH 7.0), then used to solubilize PLN (2.5 mg) by vortex. Separately, long-chain lipids were suspended into 250 μL of buffer. PLN in DHPC and long-chain lipids, both pre-chilled in ice, were combined and vortexed while allowing the sample to reach room temperature, then placed back on ice. The process was repeated at least three times to fully solubilize long-chain lipids, which produced a completely transparent liquid at cold temperature (micelle phase) and transparent solid gel at room temperature (bicelle phase). The sample was placed on ice, brought to pH 4.2 with KOH, then concentrated to ~180 μL using a 0.5 mL 10 kDa MWCO centrifugal filter (Amicon) at 4°C. The solubility of PLN^AFA^ and pPLN^AFA^ was significantly diminished at higher pH. The sample was doped with 0.8 μL of 1 M YbCl_3_, corrected back to pH 4.2 with KOH, then loaded into a 5 mm flat bottom sample cell (New Era).

For bicelles containing the SERCA/PLN^AFA^ complexes, SERCA1a was purified from rabbit skeletal muscle as previously described^4^. SERCA was eluted from Reactive Red affinity resin at ~0.5 mg/mL in SERCA, 0.1% C_12_E_8_, 1 mM CaCl_2_, 1 mM MgCl_2_, 20 mM MOPS, 20% glycerol, 8 mM ADP, 0.25 mM DTT, pH 7.00 and stored at −80°C. Protein concentration was determined by Pierce BCA Assay (Thermo Scientific) and activity confirmed by coupled-enzyme assays^3^. Immediately prior to use, SERCA (4 mg) was thawed at 4°C and combined with long-chain lipids (prepared as above, except with molar PE-DTPA chelating lipid) solubilized into 1 mL of 4% C_12_E_8_ in sample buffer. The mixture was diluted to ~30 mL with sample buffer and stirred at 4°C for 30 min prior removal of C_12_E_8_ by adding 4 g of Bio-Beads SM-2 (Bio-Rad) in stages of 0.5, 0.5, 1 and 2 g with 15 min stirring between additions. Stirring continued overnight at 4°C. Bio-Beads were removed by a 25G syringe and the cloudy suspension of proteoliposomes centrifuged at (12,000 rpm, JA25.50 rotor, 4°C, 30 min). The pellet was resuspended into ~40 mL sample buffer and centrifuged once more to wash out residual elution buffer. The final pellet was resuspended with 250 μL sample buffer and fully solubilized by adding a 250 μL mixture of PLN^AFA^, or pPLN^AFA^ (0.23 mg), in DHPC (adjusted to pH 7.0) with several cooling/heating cycles under vortex. The bicelle mixture (~1 mL) was centrifuged (13,400 rpm, Eppendorf F45-12-11 rotor, 4°C, 30 sec) to remove insoluble debris and the supernatant concentrated to ~200 μL using a 0.5 mL 10 kDa MWCO centrifugal filter (Amicon) at 4 °C. The sample was doped with 1.6 μL of 1 M YbCl_3_ in four stages, correcting pH back to 7.0 with KOH at each addition. Sample buffer included 20 mM HEPES, 100 mM KCl, 1 mM NaN_3_, 5 mM MgCl_2_, 2 mM DTT, 2.5% glycerol, pH 7.0 with 4 mM EGTA or 5 mM CaCl_2_ to stabilize *E*2 or *E*1 states, respectively. SDS-PAGE confirmed co-reconstitution of SERCA and PLN in the bicelles (Extended Data Fig 12d).

**Fig. 5:**
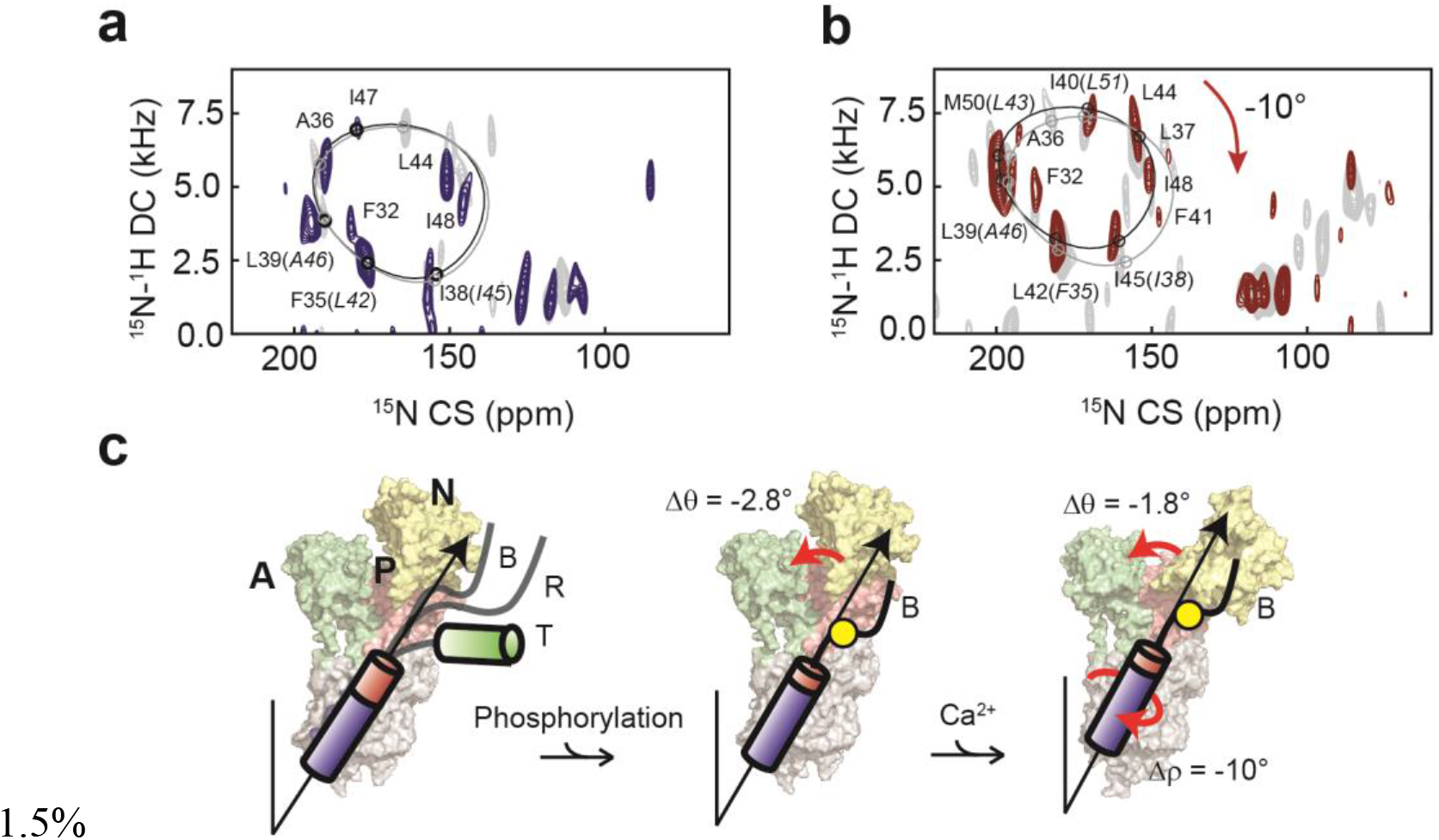
Effects of Ca^2+^ binding to SERCA on the topology of PLN and pPLN. **a, b,** 2D [^15^N-^1^H] SE-SAMPI4 spectrum of PLN^AFA^ (**a**) and pPLN^AFA^ (**b**) bound to SERCA in the *E*1 form reconstituted into aligned lipid bicelles. PISA wheels for an ideal helix [(ϕ, ψ)=(−63°, −42°)] are superimposed. Equivalent spectra of the *E*2 form complexes are shown in grey. **c,** The conformational and topological effects of phosphorylation and Ca^2+^ binding to the SERCA/PLN^AFA^ complex.

### Synthesis of TFMB and tagging of SERCA

Trifluoromethylbenzyl (TFMB)-methanethiosulfonate (MTS) was synthesized analogously to our method previously reported for synthesizing a ^13^C-ethylmethanethiosulfonate tag^5,6^ with minor modifications. Briefly, 4-(trifluoromethyl)benzyl bromide (5 mmole), MTS (5 mmole) and KI (0.03 mmole) were dissolved into 2 mL of dimethylformamide and stirred under nitrogen for 16 h at 40°C.

For TFMB tagging, SERCA (8 mg) was thawed and dialyzed into 1 L of *E*1-state sample buffer (as above, but without DDT and including 0.25 mM C_12_E_8_) overnight at 4 °C. DMPC (25.2 mg) and POPC (7.1 mg), solubilized in 500 μL of 5% C_12_E_8_, was then added to dialyzed SERCA. Detergent was removed by stirring with 1 g Bio-Beads SM-2 for 1.5 h at 4°C and a further 30 min at room temperature. Proteoliposomes were removed from Bio-Beads with a 25G syringe and centrifuged (18,000 rpm, JA25.50 rotor, 4 °C, 40 min). The pellet was suspended into 120 μL buffer and solubilized by adding DHPC (*q* = 4 for oriented or *q* = 0.5 for isotropic bicelles). TFMB-MTS (200 mM in DMSO) was added to the bicelles at 5:1 molar excess and incubated at room temperature for 1 hour prior to concentrating to ~250 with a 0.5 mL 10 kDa MWCO centrifugal filter (Amicon) and loading into a Shigemi NMR tube for ^19^F NMR measurement.

^19^F NMR spectra of TFMB-tagged SERCA were acquired on a solution state Bruker 600 MHz Avance NEO spectrometer equipped with a TCI HCN cryoprobe. 1D single-pulse experiments were acquired using a 90° pulse of 12 μs and recycle delay of 0.4 s. Spectra in isotropic bicelles were acquired with 1 k scans and 4 k scans for oriented bicelles. Spectra were processed using NMRPipe^7^.

### Oriented solid-state NMR spectroscopy

All ^15^N spectra were acquired on a Varian VNMRS spectrometer equipped with a low-*E* static bicelle probe^8^ operating at a ^1^H frequency of 700 MHz. 1D [^1^H-^15^N]-cross-polarization (CP)-based experiments used 90° pulse length of 5 μs, or 50 kHz radiofrequency (RF) field, on ^1^H and ^15^N channels; contact time of 500 μs with a 10% linear ramp on ^1^H centered at 50 kHz; and an acquisition time of 10 ms under 50 kHz SPINAL64 heteronuclear proton decoupling^9^. For all experiments, ^15^N was set to 166.3 ppm and externally referenced to ^15^NH_4_Cl at 39.3 ppm^10^; and detected using a spectral width of 100 kHz. 1D [^1^H-^15^N] water-edited (WE)-CP spectra^11^ spectra were acquired with a 1.6 ms spin-echo period following an initial 90° pulse on ^1^H as a *T*_*2*_-filter to dephase magnetization not belonging to water, then a 100 ms longitudinal mixing period to transfer ^1^H polarization from water to protein before the final Hartmann-Hahn^12^ CP spin-lock to finally transfer ^1^H magnetization to ^15^N. 1D [^1^H-^15^N] rINEPT spectra^13,14^ utilized a 2τ INEPT period of 10.4 ms and an acquisition time of 30 ms under WALTZ16 heteronuclear decoupling^15^.

2D separated local field (SLF) spectra were collected using a signal-enhanced (SE)-SAMPI4 experiment^16–18^. The indirect dipolar dimension utilized complex points and a spectral width of 20.8 kHz. The *t*_1_ evolution period utilized ^1^H homonuclear decoupling with an RF field of 50 kHz and 48 μs dwell time, and a phase-switched spin-lock pulses on ^1^H and ^15^N of 62.5 kHz RF field. The sensitivity enhancement block used a τ delay of 75 μs and three cycles of phase-modulated Lee-Goldberg (PMLG) homonuclear decoupling^19^ with an effective RF field of 80 kHz.

3D SE-SAMPI4-PDSD spectra^20^ were acquired with 15 increments in both indirect dimensions; spectral widths of 20.8 kHz and 10 kHz in the dipolar coupling and indirect ^15^N dimensions, respectively; and 3 s mixing time for ^15^N-^15^N diffusion^21^. Total acquisition times for PLN^AFA^ samples were typically 1 h for a 1D [^1^H-^15^N] CP spectra (1 k scans), 5 h for 1D [^1^H-^15^N] WE-CP spectra (5 k scans), 24 h for 1D [^1^H-^15^N] rINEPT spectra (25 k scans), 40 h for 2D SE-SAMPI4 spectra (1 k scans) and 2 weeks for 3D SE-SAMPI4-PDSD spectra (2 experiments added with 0.25 k scans each). For the SERCA/PLN^AFA^ complexes, 2D SE-SAMPI4 spectra were acquired over 4.5 days at 25 °C (4 k scans, 15 indirect points) and 1D [^1^H-^15^N] CP spectra for 4 h (4 k scans). A recycle delay of 3 sec was used for all experiments. All spectra were processed using NMRPipe^7^ and analyzed using NMRFAM-SPARKY^22^ and Nmrglue^23^.

### PISA wheel simulations and fitting

Polar Index Slant Angle (PISA) wheels^24,25^ were fitted to 2D SE-SAMPI4 spectra using the PISA-SPARKY plugin in the NMRFARM-SPARKY package^26^. Default parameters were used to describe ideal helix structure and ^15^N chemical shift (CS) and ^15^N-^1^H dipolar coupling (DC) tensors. The Cα-N-H bond angle was modified from 116° to 119° to best fit PLN^AFA^ spectra. CSs and DCs were fit by exhaustively searching tilts (θ), rotations (ρ_L31_, *i.e.*, referenced to residue Leu31) and order parameters (*S*) in increments of 0.1°, 1.0° and 0.01, respectively, for the lowest RMSD between calculated and experimental values^26^. The parameter *S*, which factors scaling due to rigid-body helical fluctuations and imperfect alignment, was determined as 0.80 ± 0.05 and 0.81 ± 0.03 for PLN^AFA^ and pPLN^AFA^, respectively, alone in bicelles. Due to the sparsity of peaks assigned for spectra of PLN in complex with SERCA, *S* was fixed to 0.80 to reduce fitting errors. Errors in tilt θ, ρ_L31_ and *S* were determined by repeating fitting 20 times with peak positions randomly adjusted at each iteration. Random adjustments were taken from a Gaussian distribution having a standard deviation equal to average FWHM peak linewidths (3 ppm for CS and 0.8 kHz for DC dimensions).

### Unrestrained molecular dynamics

A simulation of truncated PLN^AFA^ (M20 to L52; PDB 2LPF^27^) in 97 DMPC and 32 POPC, 150 mM KCl and 4927 waters was constructed using the CHARMM-GUI webserver^28,29^ and CHARMM36 forcefield^30^. Production runs were done using the AMBER18^31,32^ package at 10 Å cutoff and 8 Å force-based switching and default configuration files provided by the CHARMM-GUI webserver. For example, using the Langevin thermostat^33^ (310 K), Monte Carlo barostat^34^ (1 bar, semi-isotropic coupling) and the SHAKE algorithm^35^ for constraining hydrogens. The simulation was run for 1 μs. The final 900 ns of trajectory was used to predict ^15^N chemical shifts and ^15^N-^1^H dipolar couplings, as previously reported^36^.

### NMR-restrained refinement of the SERCA/PLN complex

The initial conformation of SERCA in the *E*2 state was obtained from the crystal structure of the SERCA/PLN complex (PDB 4Y3U)^37^. Missing loops in the structure of SERCA were introduced using MODELLER software^38^. The interface between SERCA and PLN^AFA^ (human sequence with a single Asn27Lys substitution) was then refined *in silico* by introducing information from our cross-linking data. Specifically, the transmembrane section of PLN^AFA^ was docked onto SERCA *in vacuo*, maintaining backbone positional restraints on the pump and restraining the dihedral angles of PLN^AFA^ to retain the helical structure. Docking was guided by cross-linking data^39-41^, performing short MD runs (500 ps) with an harmonic upper wall potential applied to restrain the distances between cross-linked residues to below 5 Å. 100 such runs were performed starting from different orientations of PLN^AFA^, and the resulting docked complexes were clustered according to the backbone RMSD of the proteins. The center of the most highly populated cluster was picked as the most representative structure and used to continue the modelling. The N-terminal segment of PLN^AFA^, not present in the original structure, was built as a random coil detached from SERCA. Ser16 was modelled both with and without the phosphorylation, generating two different PLN^AFA^/SERCA complexes.

The structures of the PLN^AFA^/SERCA and pPLN^AFA^/SERCA complexes were then embedded in DMPC:POPC (3:1) bilayers, mimicking the experimental conditions, and solvated with TIP3P water^42^. The systems were equilibrated for 1 μs with positional restraints using the coarse-grained model MARTINI^43^ and then re-converted to full-atom descriptions using the Backward approach^44^. The full-atomic systems were equilibrated for 50 ns at 300 K and a further 50 ns after releasing the positional restraints. Eight equally spaced structures were extracted from the final 20 ns of sampling. These eight structures were used as starting points for ssNMR-restrained replica-averaged sampling (RAOR-MD). The final equilibrated box (of dimensions 10.8 x 10.8 x 15.9 nm^3^) contains 247 DMPC lipids, 82 POPC molecules, 39,017 TIP3P waters and 23 Na^+^ ions to neutralize the system (~174,000 atoms).

^15^N chemical shift (CS) and ^15^N-^1^H dipolar coupling (DC) restraints were incorporated into the sampling using replica-averaged restrained MD, as previously described^45,46^, composed of eight replicas evolving simultaneously. The restraining forces were gradually incorporated during an initial equilibration phase of 20 ns, where the forces were linearly increased to a maximum of 50 J/(mol·ppm^2^) and 800 J/(mol·kHz^2^) for CSs and DCs, respectively.

In order to enhance the conformational sampling of the disordered domain Ia of PLN^AFA^, we implemented a sampling based on annealing cycles, whereby the PLN^AFA^ N-terminus periodically binds and detaches from the SERCA surface. The experimental PRE measurements^47^ were incorporated in order to drive each binding event. At the beginning of each cycle, the domain Ia of PLN^AFA^ is fully detached from SERCA by introducing a lower-wall harmonic potential that pushes PLN^AFA^ residues away from SERCA residues present in the cytoplasmic domains of the pump. The lower wall potential pushes PLN^AFA^ residues 0 to 10 at least 50 Å away from Ile140 (SERCA A domain), Thr430 (SERCA N domain) and Cys674 (SERCA P domain). Residues 11 to 14 were pushed at least 25 away from those residues and in addition the interfacial SERCA residues Thr742, Ala327 and Leu119. The detachment occurs gradually, by linearly increasing the force of the lower-wall potential to a maximum of 5 J/(mol · nm^2^) after 500 ps. During this time, the temperature is also linearly increased to 370 K to enhance the conformational sampling space of the disordered domain Ia. CS and DC force constants were linearly decreased to half their value to avoid instabilities during this high-energy phase of the cycle. This detachment step yields a fully detached PLN^AFA^ domain Ia with no SERCA contacts and enough surrounding free space to explore its unbound disordered conformational space. This step is followed by another 500 ps of sampling at 370 K, where the detached PLN^AFA^ domain Ia is allowed to fluctuate. Subsequently, a binding stage follows, whereby the lower-wall potential is linearly removed over 1 ns of sampling. The PRE-derived distance restraints were gradually incorporated over this time as an upper-wall potential at 30 Angstrom with a maximum force of 5 J/(mol·nm^2^). During this step the force constants of CS and DC are also restored to their full value, and the temperature is linearly decreased to 300 K. After the binding phase, the PLN^AFA^ N-terminus adopts a SERCA-bound conformation. This step leads onto the sampling phase of the cycle, where the restraint forces and the temperature are kept constant for 2 ns. The structures sampled during these 2 ns are the ones included in the final conformational ensembles of SERCA/PLN^AFA^. This annealing cycle sampling approach is illustrated in Extended Data Fig. 13.

We performed 25 annealing cycles per replica for both PLN^AFA^ and pPLN^AFA^, resulting in a total simulation time of 0.8 μs for each ensemble. 10,000 equally separated structures in the sampling part of the cycles were extracted to perform the analyses described in the main text. All samplings were performed using a previously described version of GROMACS^48^, modified to include the CS and DC restraints^46^. The CHARMM36 force field^30^ was used. Temperature was coupled using the v-rescale algorithm^49^ and pressure was coupled at 1 bar using the semi-isotropic Berendsen method^50^. All simulations were carried out under periodic boundary conditions. The integration timestep was set to 2 fs and the LINCS algorithm was used for constraints^51^. Electrostatic interactions were accounted for using the Particle Mesh Ewald method^52^. Ensembles were analyzed using CPPTraj^53^ for principle component analyses (PCA); VMD^54^ for contact, hydrogen bond and distance measurements; custom Python scripts for computing helical tilt (θ) and rotation (ρ) angles; and GROMACS energy tool for electrostatic interactions.

## Extended data figures and tables

**Extended Data Fig. 1:**
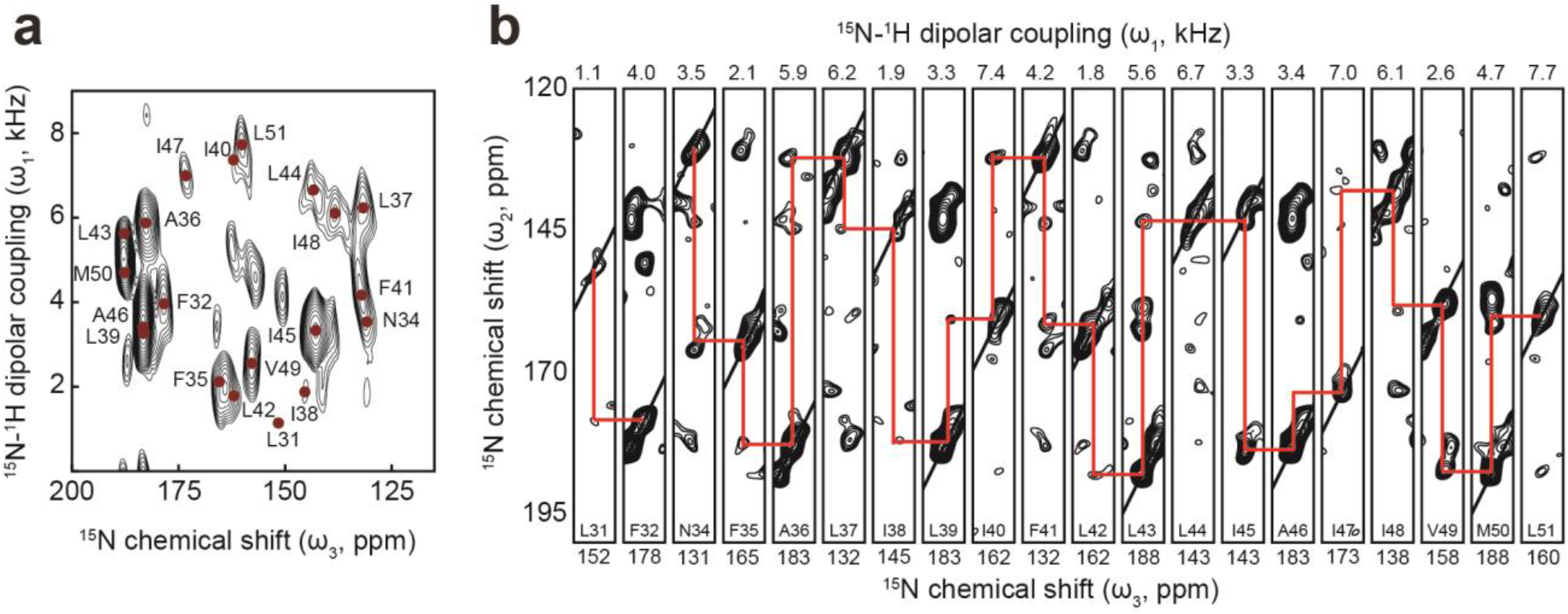
Assignment by 3D OS-ssNMR. **a,** 2D projection and **b,** strip plot of [^15^N-^15^N-^1^H] 3D SE-SAMPI4-PDSD spectrum of pPLN^AFA^ in bicelles showing sequential ^15^N-^15^N correlations.

**Extended Data Fig. 2:**
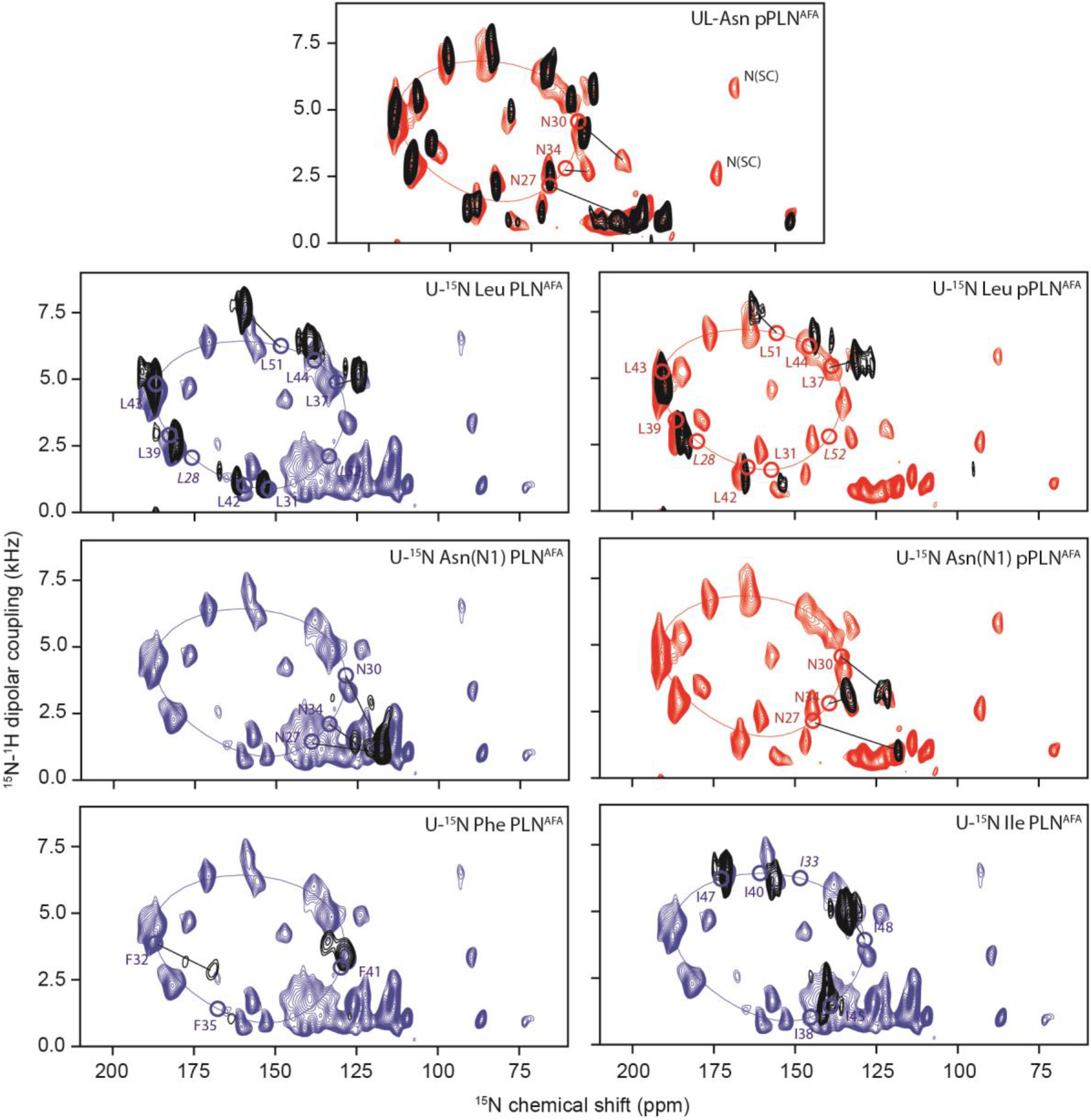
Assignment of OS-ssNMR spectra by selective labeling and un-labeling experiments. 2D [^15^N-^1^H] SE-SAMPI4 spectra of uniformly ^15^N-labelled PLN^AFA^ (blue) and pPLN^AFA^ (red) overlaid with recombinant residue-specific uniformly labeled (U) and uniformly unlabeled (UL) preparations (black). Circles indicate theoretical peak positions based on PISA fitting and labels in italics are residues missing from experimental spectra.

**Extended Data Fig. 3:**
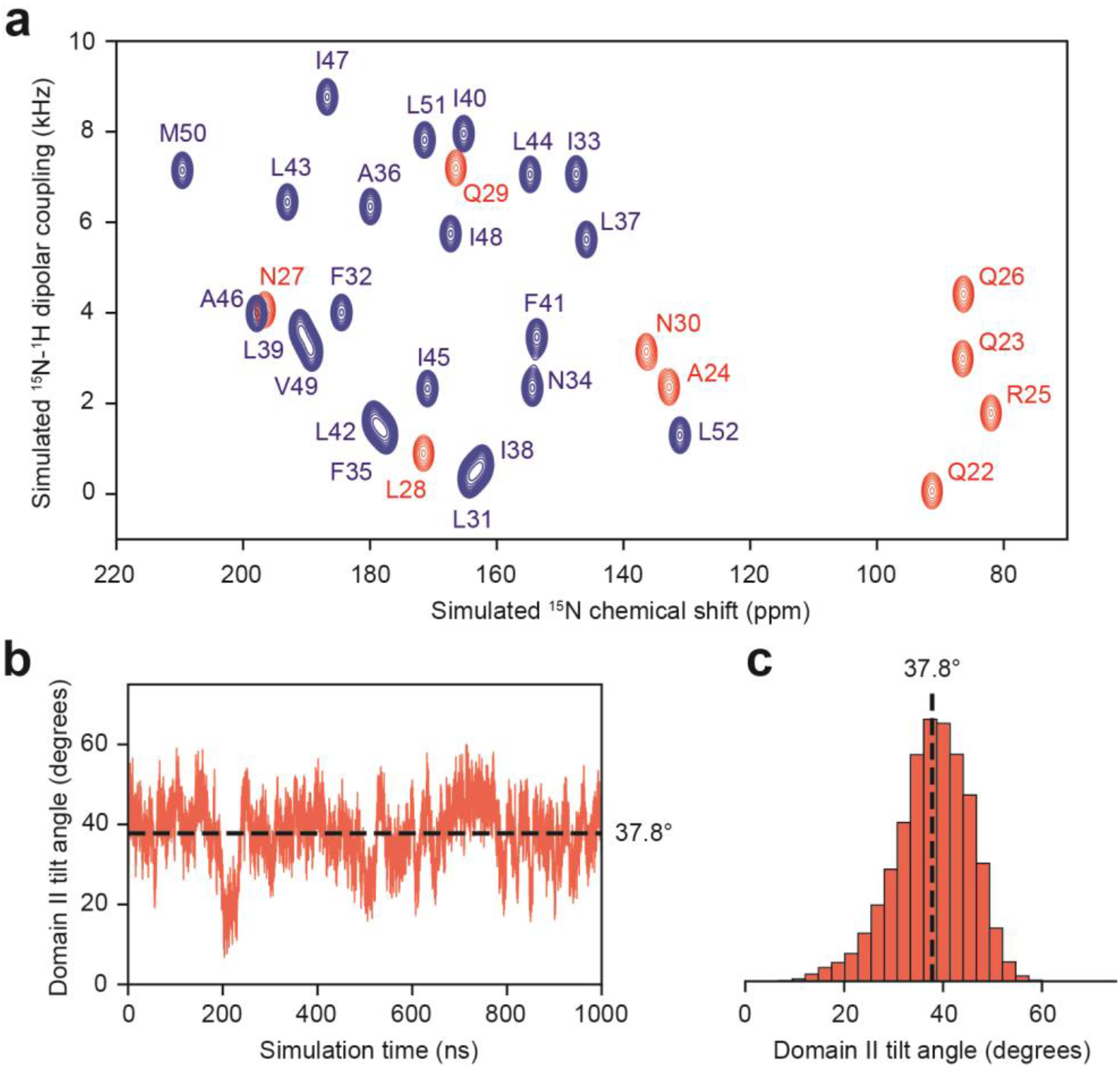
OS-ssNMR prediction of SLF spectra by unrestrained MD simulation. Unrestrained 1 μs of MD simulation of truncated PLN^AFA^ (M20 to L52; PDB 2LPF^27^) embedded in a DMPC/POPC bilayer showing **a,** the back-calculated SLF spectrum of domain Ib (red) and II (blue) residues computed as previously described^36^, **b,** PLN^AFA^ domain II tilt angle throughout the simulation and **c,** the corresponding histogram.

**Extended Data Fig. 4:**
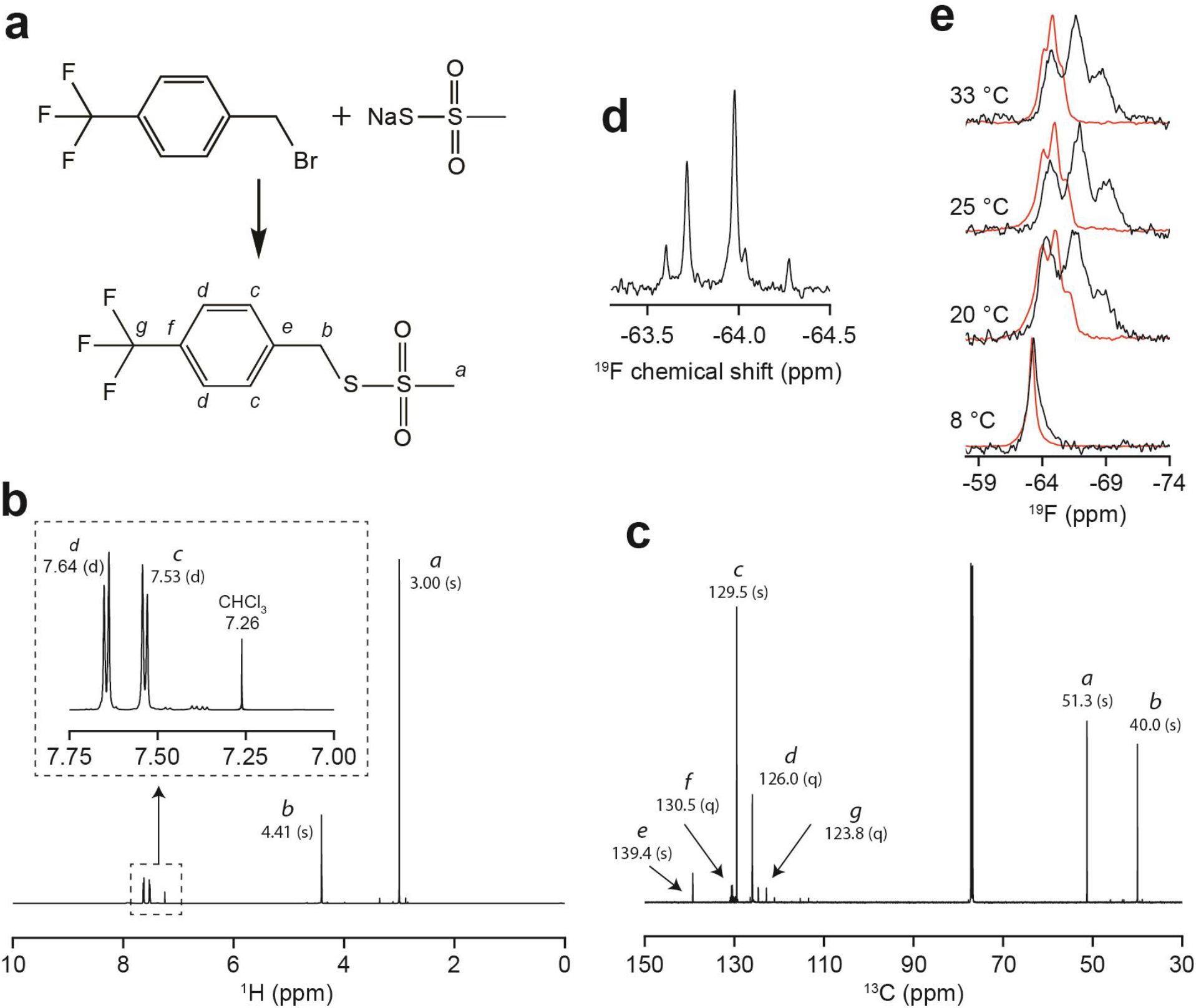
Synthesis and oriented NMR of the trifluoromethylbenzyl (TFMB)-methanethiosulfonate (MTS) tag. **a,** Synthesis of the TFMB-MTS tag. Lettering specifies carbon and proton assignments in **b,** ^1^H and **c,** ^13^C NMR spectra of the tag in CDCl3. Values in parenthesis specify peak multiplicity (s = singlet, d = doublet, q = quartet). **d,** ^19^F NMR of TFMB-tagged SERCA in isotropic bicelles. **e,** ^19^F NMR of TFMB-tagged SERCA (black) and free tag (red) in oriented bicelles. Spectra show that the tag partitions into bicelles, but with considerably reduced dipolar coupling compared to SERCA-reacted tag. This confirms that the dipolar triplet is not from unreacted tag.

**Extended Data Fig. 5:**
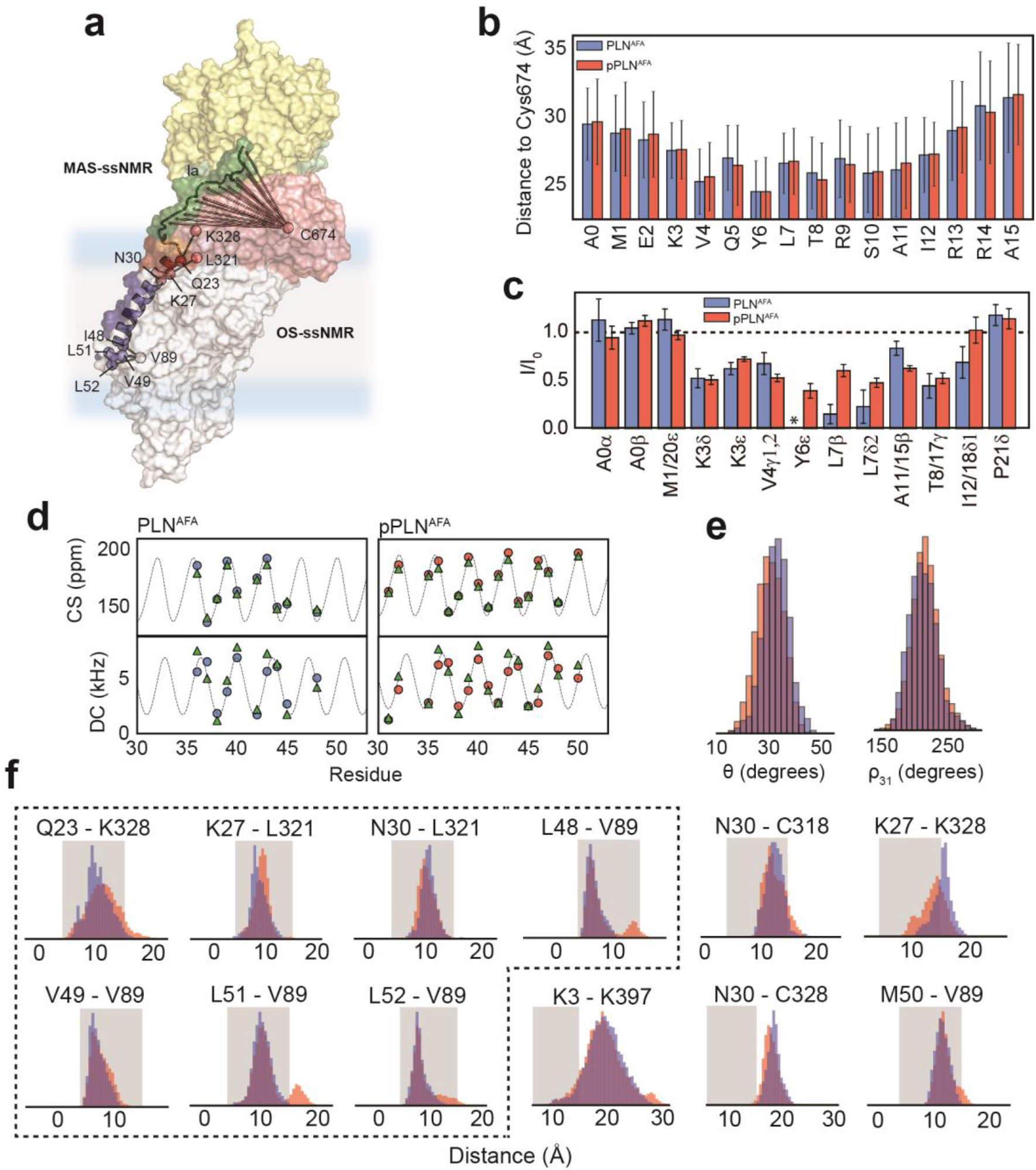
Restraints for RAOR-MD structural refinement. **a,** Schematic of crosslinking restraints^39–41^ used for docking and paramagnetic relaxation enhancements previously observed by MAS-ssNMR^47^ between methanethiosulfonate spin-labeled (MTSSL) Cys674 and cytoplasmic residues of PLN^AFA^ and pPLN^AFA^. **b,** Averaged pairwise distances between cytoplasmic residues (at Cβ) of PLN^AFA^ and Cys674-Sγ show qualitative agreement with **c,** prior [^13^C-^13^C] DARR MAS-ssNMR PRE measurements^47^. **d,** Wave plots comparing ^15^N chemical shifts and ^15^N-^1^H dipolar couplings obtained experimentally (blue or red circles) and back-calculated (green triangles) from RAOR-MD ensembles. Wave lines were taken from a PISA fits to the experimental points. **e,** Tilt (θ) and azimuthal angles (ρ31) back-calculated from RAOR-MD ensembles of PLN^AFA^ (blue) and pPLN^AFA^ (red). **f,** REMD pairwise distance distributions (Cβ-Cβ) of crosslinking residues^39–41,55,56^. Histograms encased in dotted lines relate to residue-pairs restrained during initial docking. Shaded regions show a typical 4 to 15 Å crosslinking range. The residue pair Asn30 – Cys318, outside of this range, has optimal crosslinking with longer (15 Å) reagents^55^.

**Extended Data Fig. 6:**
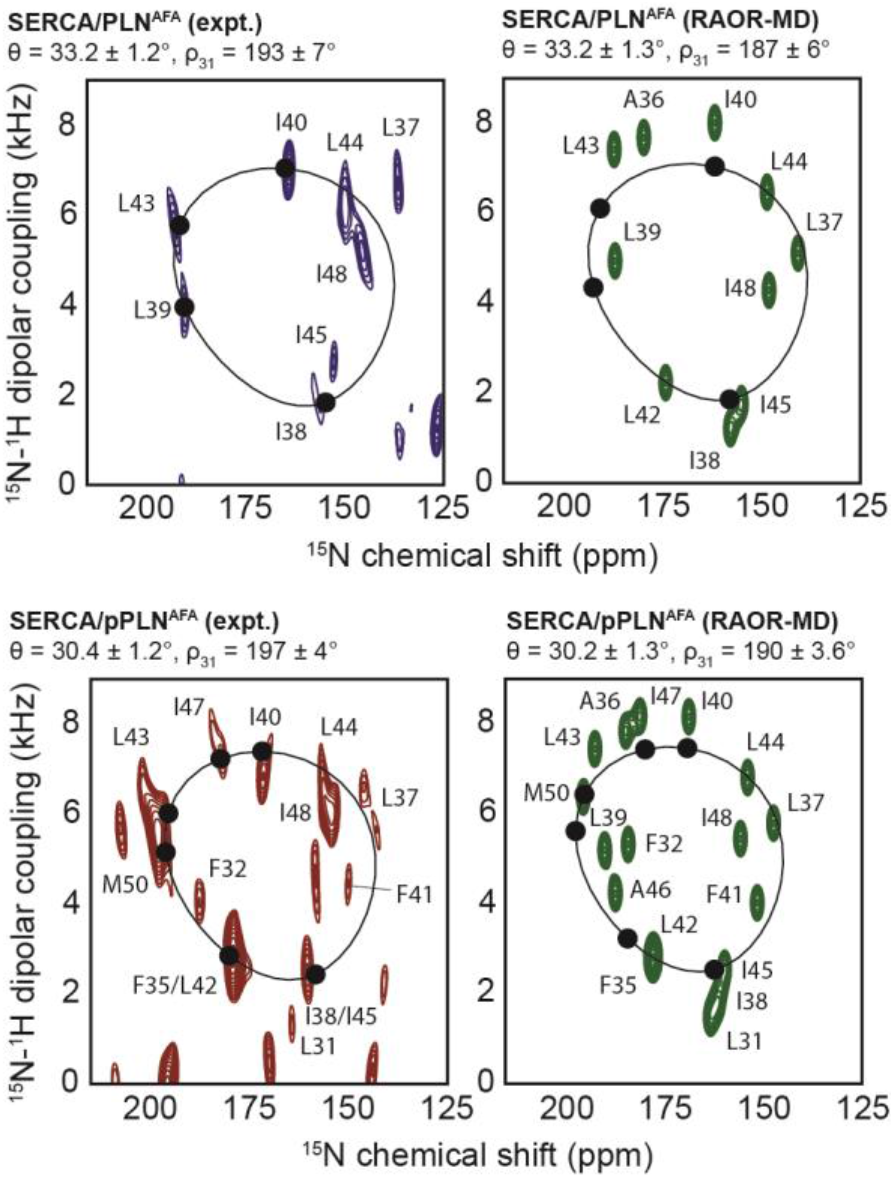
Experimental and back-calculated RAOR-MD SLF spectra of the PLN^AFA^/SERCA complex. Comparison of cross-peak positions observed from experimental SE-SAMPI4 spectra (blue or red) and back-calculated from RAOR-MD ensembles (green). PISA fits were performed using only residues (solid circles) conforming closely to ideal helicity in experimental spectra (*i.e*. Ile38, Leu39, Ile40 and Leu43 for PLN^AFA^; and Phe35, Ile38, Ile40, Leu43, Ile47 and Met50 for pPLN^AFA^).

**Extended Data Fig. 7:**
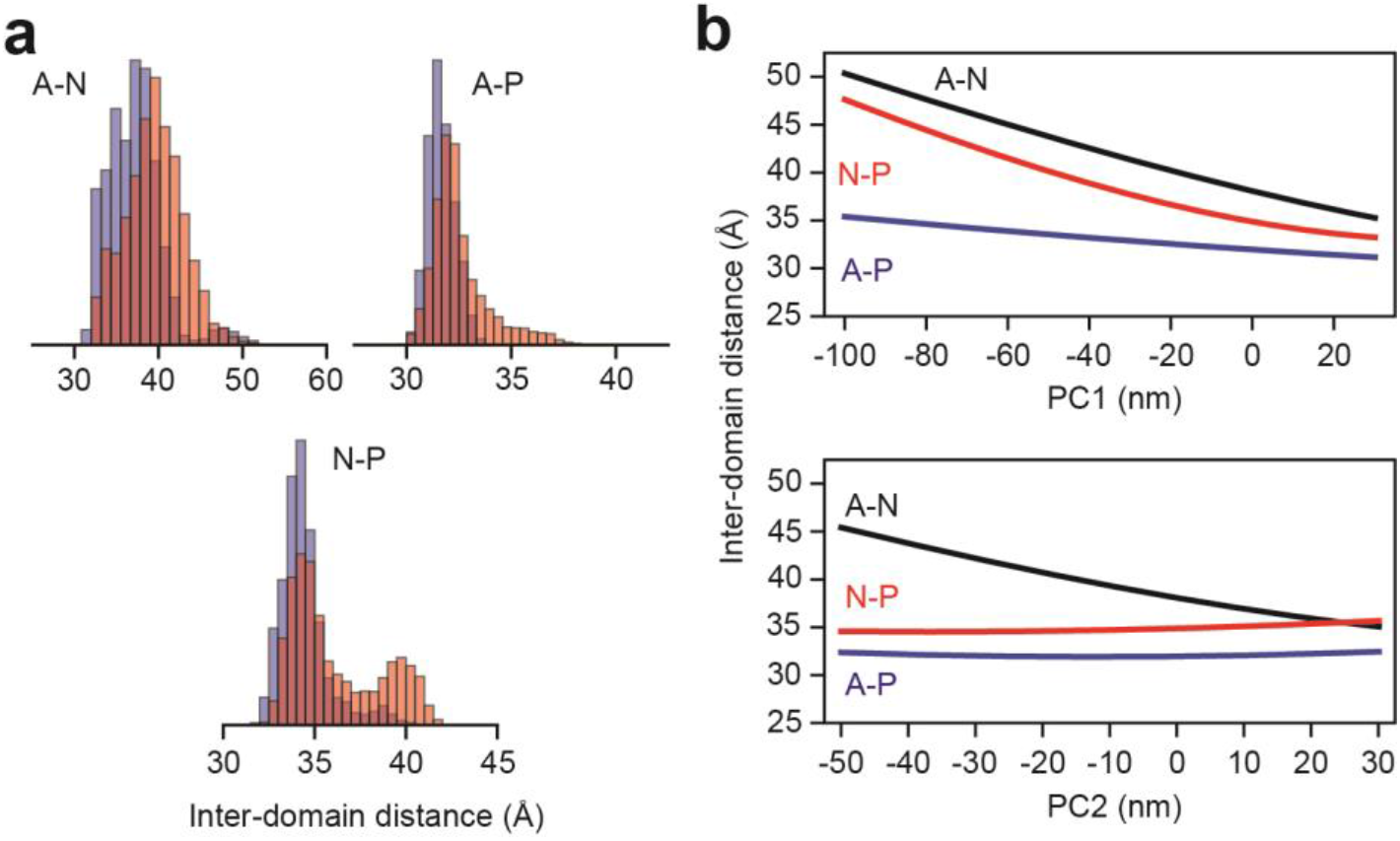
Headpiece dynamics of SERCA in RAOR-MD. **a,** Inter-domain headpiece distances distributions from RAOR-MD of SERCA in complex with PLN^AFA^ (blue) and pPLN^AFA^ (red). **b,** SERCA headpiece distance as a function of the two largest components (PC1 and PC2) from Principal Component Analysis (PCA) of the two RAOR-MD calculations. Center of masses were determined using only backbone atoms with the Actuator (A) domain defined by residues 1 to 48 and 111 to 253, the Nucleotide-binding (N) domain by 360 to 602 and the Phosphorylation (P) domain by 314 to 359 and 602 to 757.

**Extended Data Fig. 8:**
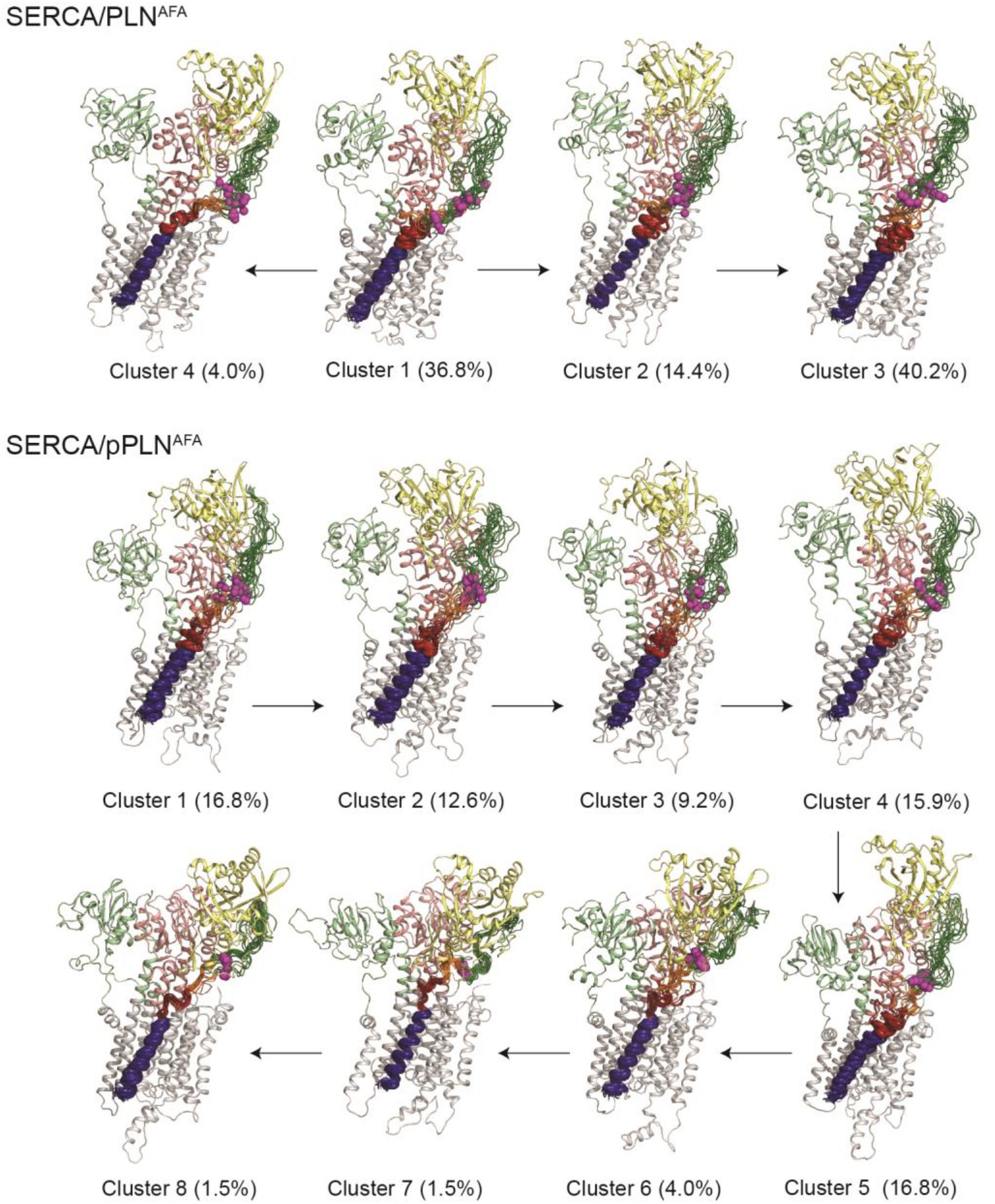
RAOR-MD clustering. Top 20 most representative structures of each cluster extracted from PCA. Structures were aligned by TM residues of SERCA. Only the most representative structure of SERCA is shown for each cluster.

**Extended Data Fig. 9:**
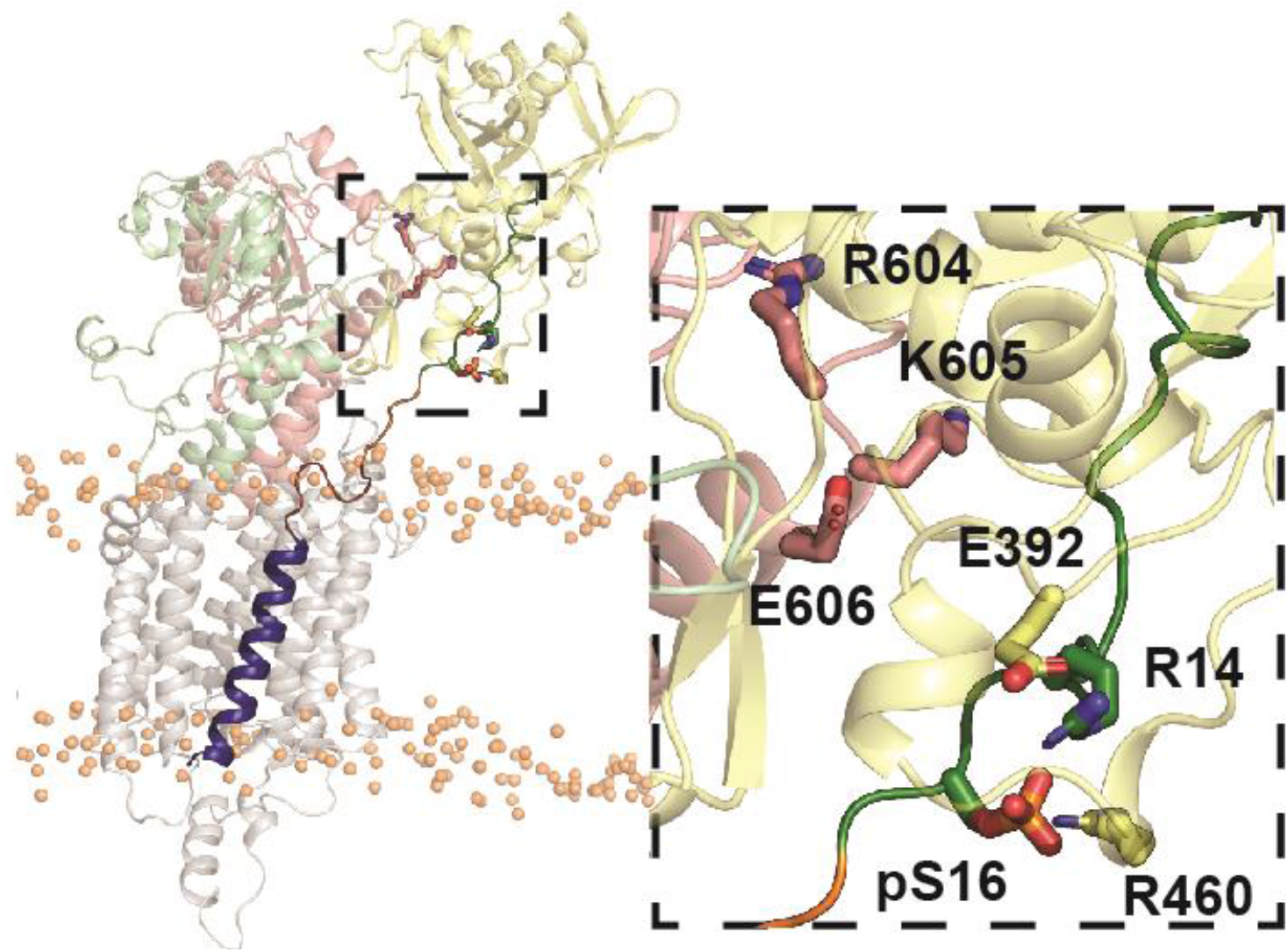
Example open state snapshot (PCA cluster 6) of SERCA stabilized by electrostatic interactions between pSer16 and the N domain.

**Extended Data Fig. 10:**
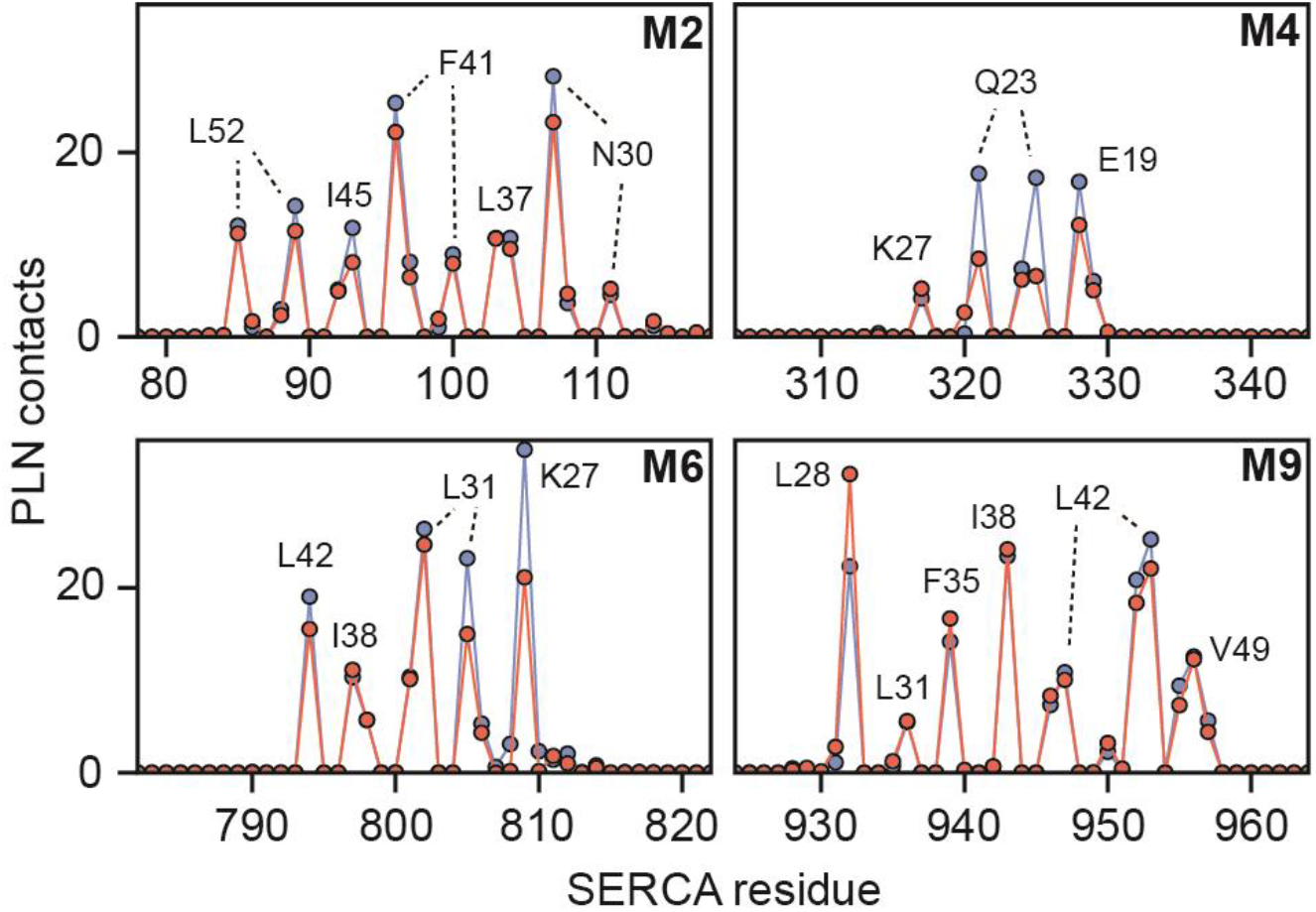
Correlation of inhibitory TM contacts with formation of the non-inhibitory B-state. **a,** Per-residue ensemble-averaged intermolecular contacts for TM helices of SERCA. A single contact by SERCA to PLN^AFA^ (blue) or pPLN^AFA^ (red) in RAOR-MD simulations is defined when any SERCA atom comes within 3.5 Å of any PLN atom for any given frame. Total contacts were accumulated by residue and the averages for each calculation was determined over 10000 frames. PLN^AFA^ residues responsible for the most contacts are designated at local maxima.

**Extended Data Fig. 11:**
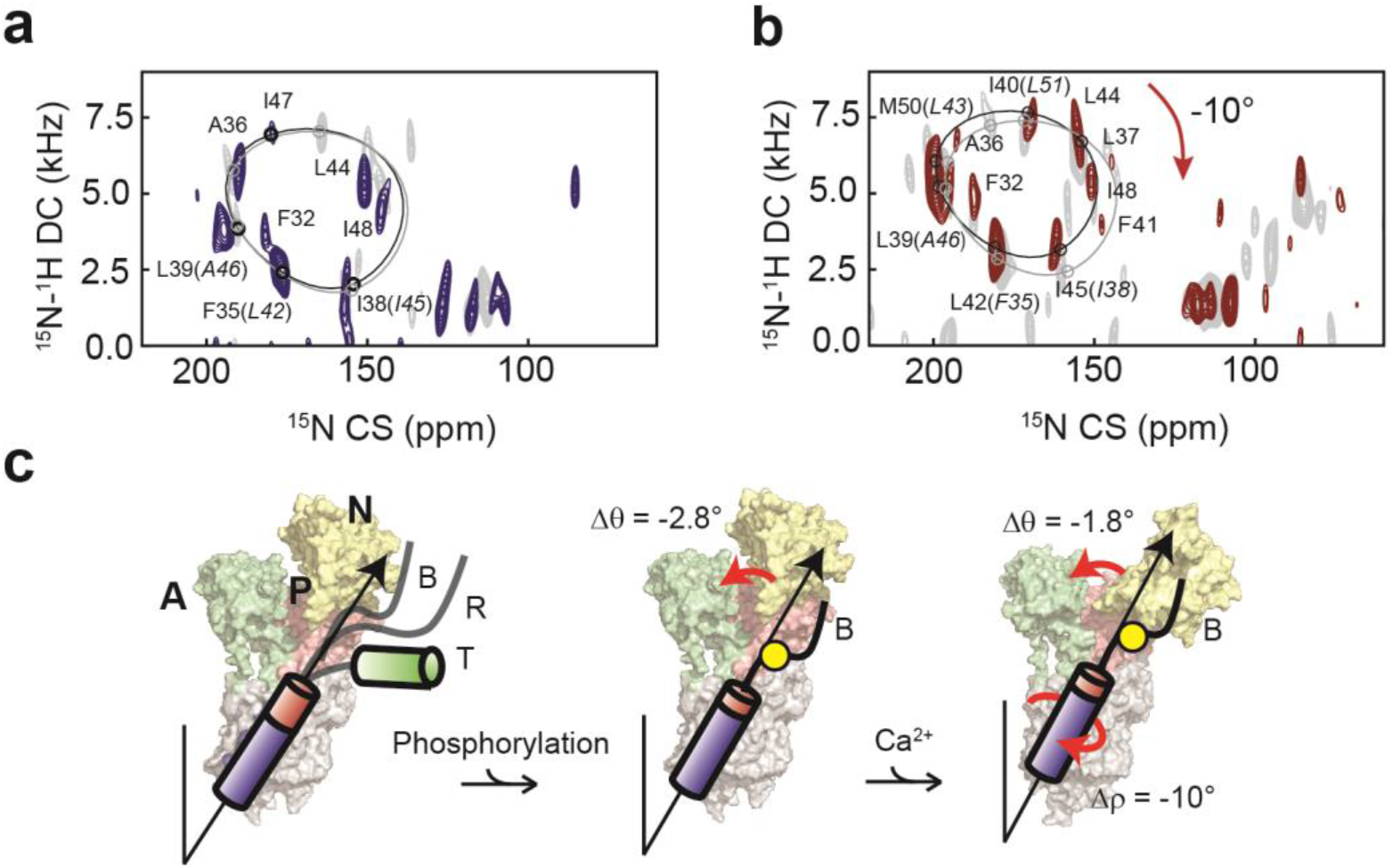
Effects of Ca^2+^ binding to SERCA on the topology of PLN and pPLN. **a, b,** 2D [^15^N-^1^H] SE-SAMPI4 spectrum of PLN^AFA^ (**a**) and pPLN^AFA^ (**b**) bound to SERCA in the *E*1 form reconstituted into aligned lipid bicelles. PISA wheels for an ideal helix [(ϕ, ψ)=(−63°, −42°)] are superimposed. Equivalent spectra of the *E*2 form complexes are shown in grey. **c,** The conformational and topological effects of phosphorylation and Ca^2+^ binding to the SERCA/PLN^AFA^ complex.

**Extended Data Fig. 12:**
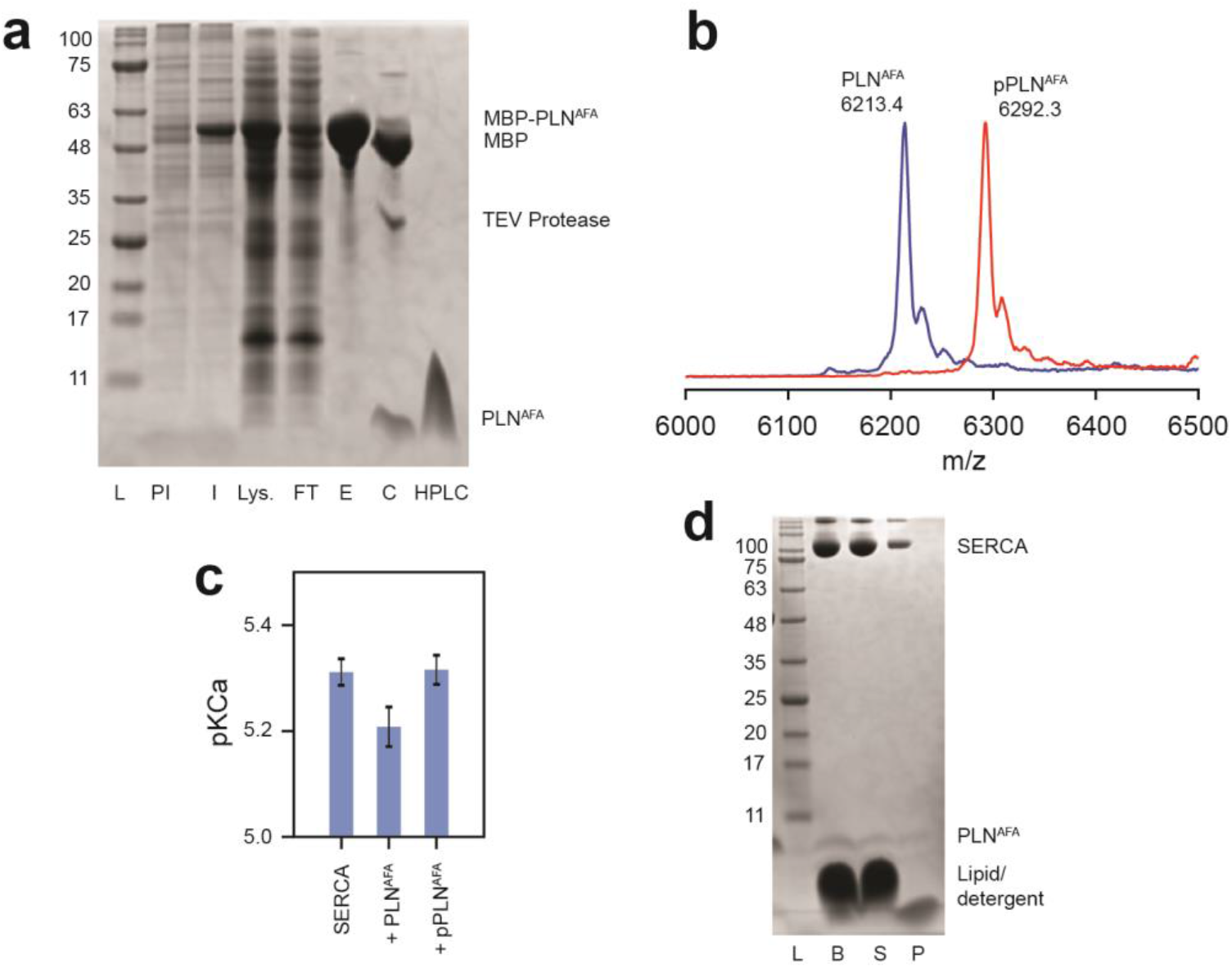
Activity and confirmation of PLN^AFA^. **a,** SDS-PAGE gel of the expression and purification of PLN^AFA^ as an MBP-fusion: ladder (L), pre-induced (PI) and induced (I) expression, lysate supernatant (Lys.), purified MBP-PLN^AFA^ eluted from amylose resin (E), MBP-PLN^AFA^ after cleavage with TEV protease (C),and HPLC-purified PLN^AFA^. **b,** MALDI-MS spectra of HPLC-purified ^15^N PLN^AFA^ and pPLN^AFA^. **c,** Coupled-enzyme ATPase activity assays of SERCA reconstituted into DMPC/POPC (4:1; lipid-to-protein ratio of 700:1) liposomes with a 5-fold excess of either PLN^AFA^ or pPLN^AFA^. Assays consisted of 50 mM HEPES, 100 mM KCl, 5 mM MgCl2, 0.2 mM NADH, 0.5 mM PEP, 10 U/mL pyruvate kinase, 10 U/mL lactate dehydrogenase, and 7 μM calcium ionophore A23187 at pH 7.0. ATPase activity was measured by the rate of reduction in NADH absorbance at 340 nm at EGTA-buffered calcium concentrations between 10^−7^ and 10^−4^ M, initiated by addition of 0.5 mM ATP. Activity as a function of calcium concentration was fitted to a Hill function to extract the pKCa values (calcium concentration at half Vmax) shown. Errors indicate the standard deviation from 3 replicate measurements. **d,** SDS-PAGE gel of SERCA and PLN reconstituted into bicelles for OS-ssNMR: bicelle sample freshly solubilized by DHPC (B), supernatant after centrifugation (S) and pellet formed from a small non-solubilized component (P).

**Fig. 13:**
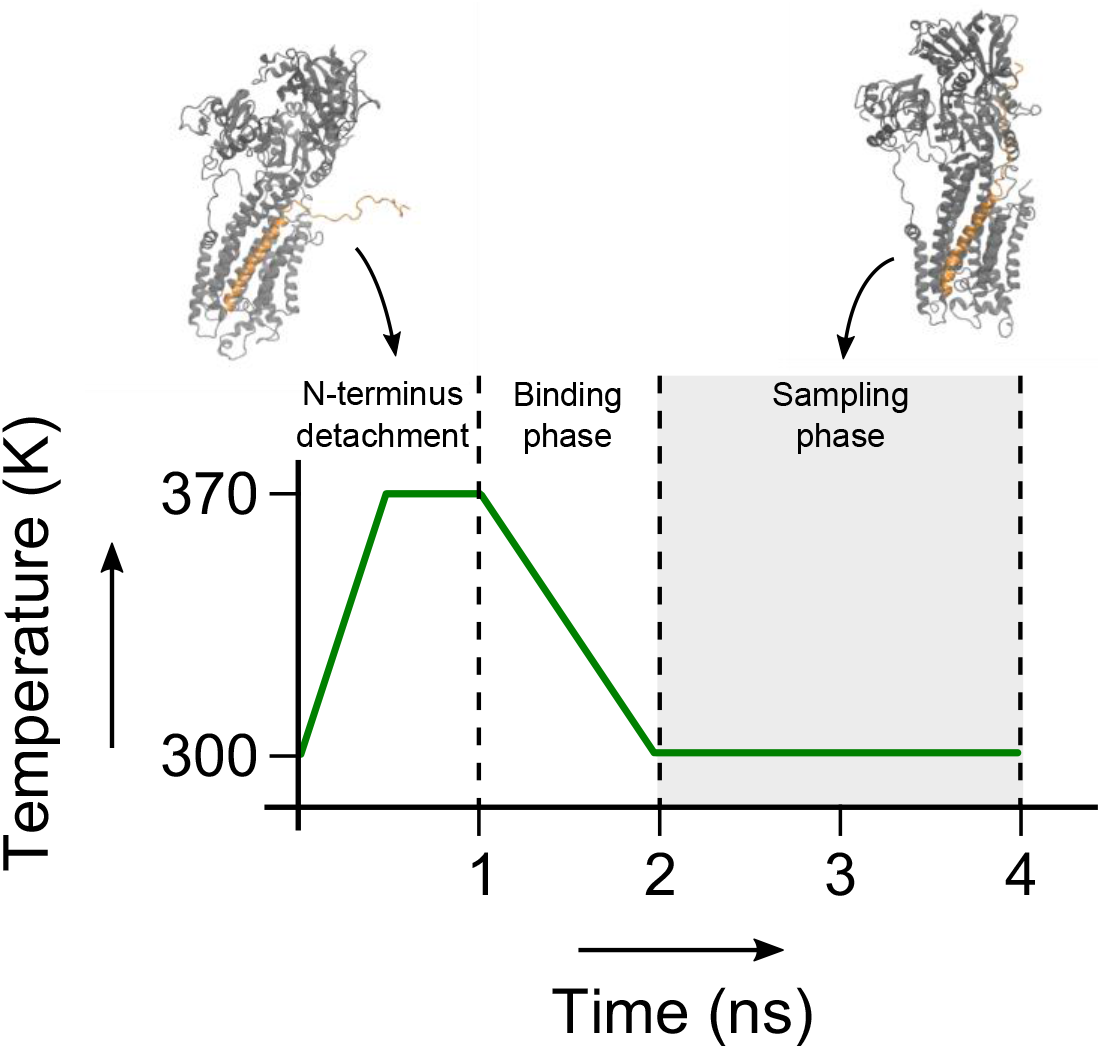
Annealing cycle used in the sampling of the SERCA/PLN^AFA^ complex. During the first nanosecond of the simulation, the N-terminus of PLN is detached from SERCA and the temperature is raised to 370 K to randomize its conformation. Then the system is cooled back down to 300 K over 1 ns, during which time the CS, DC and PRE restraints are re-introduced. The complex is sampled for 2 ns, after which the cycle restarts.

**Table 1:** OS-ssNMR assignments. Summarized ^15^N CS and ^15^N-^1^H DC assignments of PLN^AFA^ and pPLN^AFA^ reconstituted alone in bicelles or in complex with SERCA in either *E*2 (Ca^2+^-free) or *E*1 (5 mM Ca^2+^) states.

| Res. | PLN<sup>AFA</sup> |  | pSer16-PLN<sup>AFA</sup> |  | PLN<sup>AFA</sup> +<br>SERCA (E2) |  | pSer16-PLN<sup>AFA</sup><br>+ SERCA (E2) |  | PLN<sup>AFA</sup> +<br>SERCA (E1) |  | pSer16-PLN<sup>AFA</sup><br>+ SERCA (E1) |  |
| --- | --- | --- | --- | --- | --- | --- | --- | --- | --- | --- | --- | --- |
|  | CS | DC | CS | DC | CS | DC | CS | DC | CS | DC | CS | DC |
| N27 | 117.6* | 1.19* | 118.1 | 0.90 | - | - | - | - | - | - | - | - |
| N30 | 117.6* | 1.19* | 121.5 | 3.10 | - | - | - | - | - | - | - | - |
| L31 | 152.8 | 0.94 | 155.2 | 0.74 | - | - | 164.3 | 1.26 | - | - | - | - |
| F32 | 167.7 | 2.46 | 177.9 | 3.38 | - | - | 187.6 | 4.07 | 181.6 | 3.78 | 186.6 | 4.56 |
| N34 | 125.8 | 1.36 | 132.4 | 2.71 | - | - | - | - | - | - | - | - |
| F35 | 164.7 | 1.17 | 166.2* | 1.36* | - | - | 179.1 | 2.86 | 176.4 | 2.30 | - | - |
| A36 | 176.5 | 4.74 | 184.3 | 5.28 | 186.5 | 5.72 | 191.0 | 6.37 | 190.4 | 5.91 | 192.8 | 6.76 |
| L37 | 123.9 | 4.89 | 132.2 | 5.71 | 136.5 | 6.66 | 146.0 | 6.57 | 146.1 | 4.73 | 151.7 | 5.87 |
| I38 | 140.8 | 1.03 | 146.5 | 1.40 | 156.9 | 1.86 | 160.3 | 2.54 | 156.4 | 2.11 | 164.9 | 2.14 |
| L39 | 181.7* | 2.31* | 185.7* | 2.86* | 190.3 | 3.88 | 194.4 | 3.99 | 194.5 | 3.76 | 198.5 | 4.75 |
| I40 | 155.7 | 6.19 | 163.5* | 6.88* | 164.0 | 7.04 | 171.5 | 6.87 | 166.7 | 6.99 | 169.9 | 7.25 |
| F41 | 127.8 | 3.39 | 134.6 | 4.19 | - | - | 150.0 | 4.46 | - | - | 148.4 | 4.33 |
| L42 | 160.5 | 0.91 | 166.2* | 1.36* | 175.5 | 1.74 | 179.1 | 2.86 | - | - | 181.1 | 2.63 |
| L43 | 187.5* | 4.17* | 190.9* | 4.46* | 192.8 | 5.77 | 198.3 | 5.76 | 196.5 | 5.89 | - | - |
| L44 | 138.0 | 6.02 | 145.6 | 6.19 | 149.9 | 6.22 | 155.5 | 6.26 | 156.0 | 5.19 | 155.1 | 6.65 |
| I45 | 139.2 | 1.80 | 144.5 | 2.64 | 152.7 | 2.74 | 160.3 | 2.54 | - | - | 162.1 | 3.57 |
| A46 | 181.7* | 2.31* | 185.7* | 2.86* | - | - | 191.4 | 2.84 | - | - | - | - |
| I47 | 170.5 | 6.35 | 175.9 | 6.65 | - | - | 181.8 | 7.23 | 179.5 | 6.74 | 181.9 | 7.99 |
| I48 | 132.7 | 4.88 | 139.2 | 5.39 | 145.5 | 5.16 | 154.3 | 6.03 | 151.2 | 4.94 | - | - |
| V49 | 157.1 | 1.75 | 160.1 | 2.26 | - | - | - | - | - | - | - | - |
| M50 | 187.5* | 4.17* | 190.9* | 4.46* | - | - | 198.0 | 5.14 | 202.5 | 4.80 | 199.9 | 5.94 |
| L51 | 158.3 | 7.06 | 163.5* | 6.88* | - | - | - | - | - | - | - | - |

\* These peaks are overlapped.

**Table 2:** Pairwise inter- and intramolecular hydrogen bond summary for PLN^AFA^/SERCA REMD PCA clusters. Parentheses report the average number hydrogen bonds to a charged residue of SERCA or PLN (bold font) observed over all frames assigned to the respective clusters. Hydrogen bonds were defined with a donor -acceptor distance less than 3 Å and a donor-H-acceptor angle less than 20°.

| Cluster | Average hydrogen bonds per frame |  |  |  |  |  |  |
| --- | --- | --- | --- | --- | --- | --- | --- |
|  | E2 | K3 | R9 | R13 | R14 | S16 | E19 |
| 1 | R556 (0.10)<br>K365 (0.08)<br>R638 (0.02) | - | E606 (0.05) | **E19 (0.01)** | E606 (0.01) | - | K328 (0.08)<br>**R13 (0.01)** |
| 2 | R556 (0.10)<br>K365 (0.04) | E644 (0.02)<br>D557 (0.02) | D557 (0.01) | E606 (0.05)<br>**E19 (0.04)** | E606 (0.02) | - | K328 (0.38)<br>**R13 (0.04)** |
| 3 | K365 (0.11)<br>R556 (0.09)<br>R638 (0.01) | D399 (0.01) | E606 (0.02) | - | E606 (0.06) | - | K328 (0.15)<br>R325 (0.04)<br>K329 (0.03) |
| 4 | K397 (0.14)<br>K365 (0.11)<br>R638 (0.05) | D399 (0.24) | D557 (0.10)<br>E392 (0.02)<br>E606 (0.01) | D616 (0.06) | - | - | - |

**Table 3:** Pairwise inter- and intramolecular hydrogen bond summary for pSer16-PLN^AFA^/SERCA REMD PCA clusters. Hydrogen bonds were measured according to Table 2.

| Cluster | Average hydrogen bonds per frame |  |  |  |  |  |  |
| --- | --- | --- | --- | --- | --- | --- | --- |
|  | E2 | K3 | R9 | R13 | R14 | S16 | E19 |
| 1 | K365 (0.04)<br>R556 (0.03)<br>K397 (0.02) | D557 (0.01) | D557 (0.14)<br>**S16 (0.13)**<br>E392 (0.02) | **S16 (0.59)**<br>E606 (0.02) | **S16 (0.72)**<br>E606 (0.04) | **R14 (0.72)**<br>**R13 (0.59)**<br>R604 (0.54)<br>K605 (0.30) | K328 (0.07) |
|  |  |  |  |  |  | **R9 (0.13)** | K329 (0.02) |
| 2 | K365 (0.07)<br>R556 (0.03) | D399 (0.01) | **S16 (0.55)**<br>E606 (0.03) | **S16 (0.21)**<br>**E19 (0.06)**<br>E748 (0.02)<br>D616 (0.01) | **S16 (0.81)**<br>E606 (0.06) | **R14 (0.81)**<br>**R9 (0.55)**<br>**R13 (0.21)**<br>K605 (0.16)<br>R604 (0.01) | K328 (0.10)<br>**R13 (0.06)** |
| 3 | K365 (0.12)<br>**K3 (0.02)**<br>R556 (0.01) | D399 (0.03)<br>D557 (0.02)<br>**E2 (0.02)** | **S16 (0.82)**<br>E606 (0.05) | **S16 (0.07)**<br>E748 (0.03) | **S16 (0.85)**<br>E606 (0.01) | **R14 (0.85)**<br>**R9 (0.82)**<br>**R13 (0.07)**<br>K605 (0.05) | - |
| 4 | R556 (0.09)<br>K365 (0.06)<br>R560 (0.02)<br>R638 (0.01)<br>K400 (0.01) | D557 (0.03) | **S16 (0.28)**<br>E606 (0.05) | **S16 (0.57)**<br>**E19 (0.03)** | **S16 (0.86)**<br>E606 (0.02) | **R14 (0.86)**<br>**R13 (0.57)**<br>**R9 (0.28)** | K328 (0.09)<br>**R13 (0.03)**<br>K329 (0.02) |
| 5 | R556 (0.07)<br>K365 (0.05)<br>**K3 (0.03)**<br>K400 (0.03)<br>R638 (0.03) | 399 (0.04)<br>**E2 (0.03)** | **S16 (0.16)**<br>E606 (0.05) | **S16 (0.77)** | **S16 (0.77)** | **R13 (0.77)**<br>**R14 (0.77)**<br>**R9 (0.16)**<br>K328 (0.03)<br>K329 (0.02) | R325 (0.19)<br>K328 (0.13)<br>K464 (0.02) |
| 6 | R556 (0.06)<br>**K3 (0.04)** | **E2 (0.04)** | D399 (0.08)<br>E394 (0.05) | **S16 (0.84)** | **S16 (1.14)**<br>E392 (0.25) | **R14 (1.14)**<br>R460 (0.89)<br>**R13 (0.84)**<br>K464 (0.11)<br>K329 (0.07)<br>R325 (0.01) | K328 (0.21)<br>K329 (0.04)<br>K464 (0.03) |
| 7 | K365 (0.14) | - | **E19 (0.03)**<br>D616 (0.02) | E748 (0.54)<br>E606 (0.03) | **S16 (0.36)** | R325 (1.18)<br>**R14 (0.36)**<br>K329 (0.31)<br>K328 (0.25) | **R9 (0.03)** |
| 8 | K397 (0.10)<br>K400 (0.03) | E644 (0.01) | E394 (0.23) | **S16 (0.67)** | **S16 (1.22)**<br>E392 (0.65) | **R14 (1.22)**<br>**R13 (0.67)**<br>R460 (1.35) | R325 (0.45)<br>K328 (0.44)<br>K329 (0.37) |

## Notes

### Competing Interest Statement

The authors have declared no competing interest.

